# A deep-time landscape of plant *cis*-regulatory sequence evolution

**DOI:** 10.1101/2025.09.17.676453

**Authors:** Kirk R. Amundson, Anat Hendelman, Danielle Ciren, Hailong Yang, Amber E. de Neve, Shai Tal, Adar Sulema, David Jackson, Madelaine E. Bartlett, Zachary B. Lippman, Idan Efroni

**Author notes:** These authors contributed equally to this work (Kirk R. Amundson, Anat Hendelman).

## Abstract

Developmental gene function is often conserved over deep time, but cis-regulatory sequence conservation is difficult to identify. Rapid sequence turnover, paleopolyploidy, structural variation, and limited phylogenomic sampling have impeded conserved non-coding sequence (CNS) discovery. Using Conservatory, an algorithm that leverages microsynteny and iterative alignments to map CNS-gene associations over evolution, we uncovered ∼2.3 million CNSs, including over 3,000 predating angiosperms, from 284 plant species spanning 300 million years of diversification. Ancient CNSs were enriched near developmental regulators, and mutating CNSs near *HOMEOBOX* genes produced strong phenotypes. Tracing CNS evolution uncovered key principles: CNS spacing varies, but order is conserved; genomic rearrangements form new CNS-gene associations; and ancient CNSs are preferentially retained among paralogs, but are often lost as cohorts or evolve into lineage-specific CNSs.

**One Sentence Summary:** Conservatory maps ancient cis-regulatory elements and uncovers regulatory evolution dynamics.

## Main Text

Cis*-*regulatory sequence conservation and divergence are central in phenotypic evolution (*1–3*). Defining principles that govern cis*-*regulatory evolution is key to deciphering the genetic basis of natural and artificial selection, and essential for targeted manipulation of gene regulation and organismal phenotypes (*4–6*). A long-standing paradox in the evolution of regulatory sequences is that homologous genes in distantly related species apparently lack cis-regulatory sequence conservation, despite similar expression patterns and functions (*7*). However, identifying cis*-*regulatory elements (CREs) and establishing their orthology across species is often technically challenging, especially over deep time.

Evolutionary conservation can reveal CREs because conservation can indicate selection for preserved function (*8–14*). Conserved non-coding sequences (CNSs) are often identified through whole-genome alignment approaches (*15*, *16*), but these methods fail in comparing distantly related species, because regulatory sequence evolves rapidly. Additionally, frequent macro- and micro-genomic rearrangements, gene duplications, and ancestral whole-genome duplications obscure orthology relationships. Thus, homologous CREs often cannot be directly aligned in distantly related species (*17*, *18*). Iterative alignment through intermediate “bridge” genomes improves CNS discovery across broader evolutionary distances (*18*, *19*). However, these strategies must still contend with pervasive structural variation and the challenges of complex, highly rearranged, paleopolyploid, or repetitive genomes typical of many eukaryotes (*20*).

The challenges of identifying CNSs are magnified in plant genomes. Repeated cycles of whole-genome duplication, subsequent fractionation to diploidization, and the unequal, often biased retention of paralogous genes obscure gene and CNS orthology relationships. These extreme patterns of genome evolution are incompatible with current whole-genome alignment tools, requiring the development of new approaches (*20*). Indeed, most identified plant CNSs to date are limited in phylogenetic scope, with only a handful extending beyond taxonomic families (*21–25*).

Here, we present Conservatory (www.conservatorycns.com), a gene orthogroup-centric CNS discovery algorithm developed for detecting noncoding sequence conservation while accounting for complex orthology relationships, extreme structural variation, and rapid sequence divergence. Using Conservatory, we annotated and resolved CNS orthologies across 314 plant genomes representing 284 species spanning green plant diversity. Our analysis uncovered widespread conservation of noncoding sequences at unprecedented evolutionary depths. We use this comprehensive atlas to trace ∼300 million years of cis-regulatory sequence evolution in plants (*26*), uncovering fundamental principles shaping the dynamic formation, conservation, and turnover of plant regulatory sequences.

### Ancient cis*-*regulatory elements are common in plant genomes

To address challenges in identifying CNSs, we developed ‘Conservatory’, a large-scale comparative genomics algorithm that captures sequence conservation across deep time, while accounting for gene duplication and rapid sequence divergence. Conservatory identifies orthologs within a particular taxonomic family, then aligns up to 120 kb of intergenic sequence surrounding each ortholog member of the orthogroup to a common reference genome. To account for frequent paleopolyploidy, fractionation, and structural rearrangements, up to 16 co-orthologs per genome are retained, with paralogs treated as independent co-orthologs. Following alignments of the regulatory regions within each orthogroup, CNSs are identified using *phyloP* (*27*), and ancestral sequence reconstruction (*FastML*) (*28*) used to infer CNS sequences at the crown nodes of each taxonomic family. Ancestral sequences are then used to search the regulatory regions of all putative orthologs outside of the taxonomic family using high-sensitivity alignment. This two-step alignment, first within and then between families, allows for CNS discovery despite divergence of flanking sequence. To account for reference genome bias, independent CNS datasets were generated using multiple reference genomes. The algorithm then uses ‘bridge genomes’ to infer the homology of CNSs that cannot be directly aligned, but can be linked by intermediate sequences (*19*). After filtering, Conservatory generates a catalog of CNSs and their orthologous relations (**fig. S1**).

We applied Conservatory to 314 genomes, representing 284 species spanning green plant diversity, using 10 reference genomes from species in the Solanaceae, Asteraceae, Brassicaceae, Fabaceae, Poaceae, and Lemnaceae families (**Fig. 1A; table S1**). We discovered 32.8 million CNSs across all genomes, and grouped them into 2.3 million unique CNSs based on sequence orthology (*29*). Conservatory CNSs were short (median length 12-40 bp; **Fig. 1B, fig. S2A; table S2**), consistent with previous reports (*21–24*, *30*, *31*). To estimate the rate of false positives resulting from annotation errors, we searched CNSs for known peptide sequences. The rate of CNS mapping to known peptides was 0.05%-1.8% (**fig. S1E**), and these sequences were removed from downstream analysis.

**Fig. 1.**
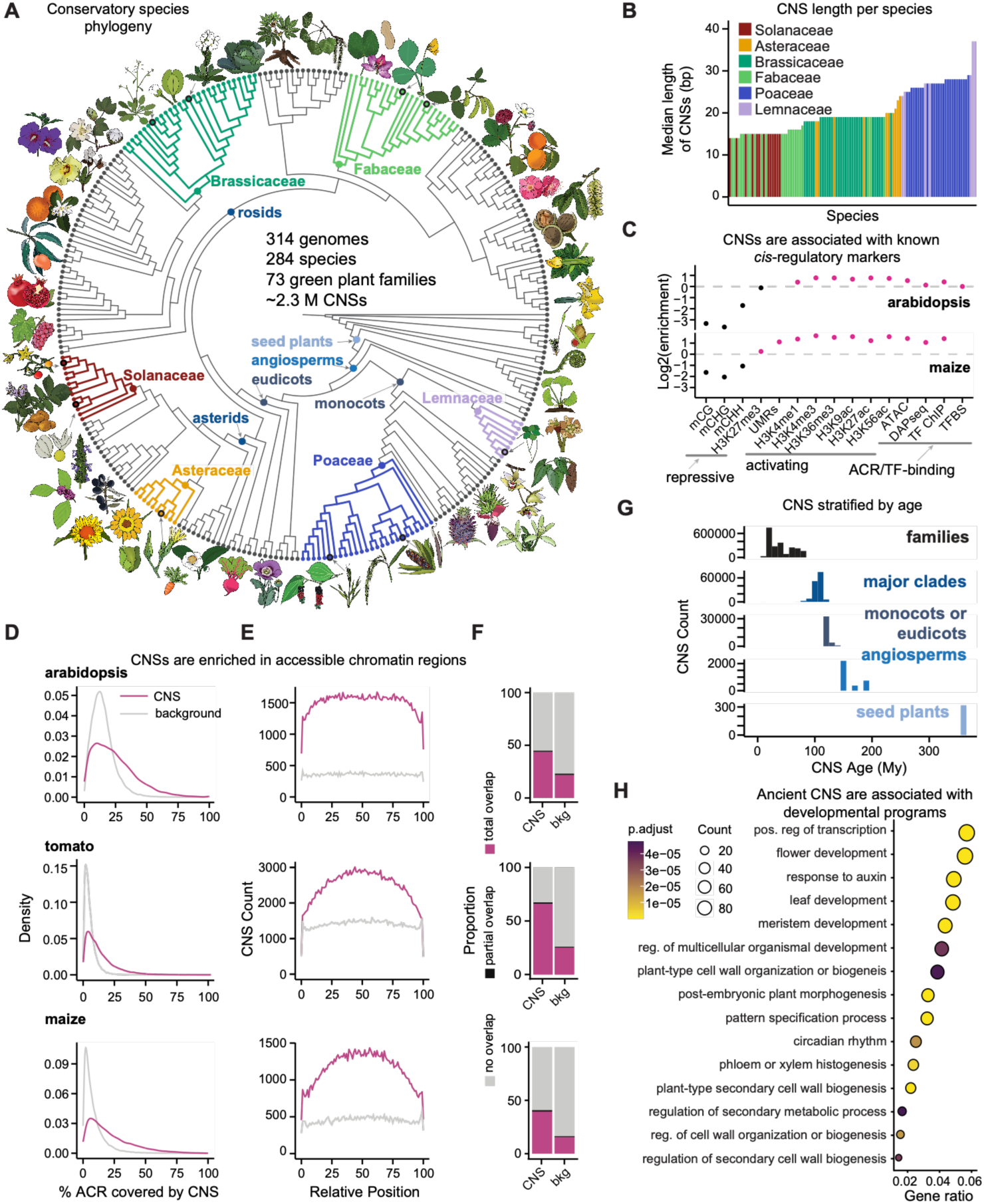
Identification of deeply conserved plant *cis*-regulatory elements. (**A**) Phylogeny of the species used in this study. The six families with reference species are colored. (**B**) Median length of CNSs, aggregated by species (table S2). (**C**) Enrichment of DNA methylation, histone modification, open chromatin, and transcription factor binding sites in CNSs, compared to intergenic background regions in Arabidopsis and maize (see **table S4** for statistics). (**D**) Distribution of the percentages of accessible chromatin regions (ACRs) of Arabidopsis, tomato, and maize that overlap CNSs (pink) versus control intergenic regions (gray)(see **table S5** for statistics). (**E**) Position distribution within overlapping ACRs for CNSs (pink) or control intergenic regions (gray). CNS are enriched at ACR midpoints (see **table S5** for statistics). (**F**) Percentage of CNSs vs control background (bkg) sequences that overlap ACRs (see **table S5** for statistics). (**G**) Frequency histograms of CNS age estimations (**table S6**). (**H**) Functional enrichment of Arabidopsis genes associated with angiosperm and seed plant CNSs. Gene ratio is the number of genes containing one or more angiosperm or seed plant CNSs that fall into the given GO category, divided by the total number of genes with angiosperm or seed plant CNSs.

Across our deep phylogenomic sampling, CNSs were enriched for transcription factor binding sites and activating histone modifications, and depleted for DNA methylation and repressive histone marks, relative to stringently-defined background regions in the Conservatory search space, as observed for other plant regulatory sequences (*22*, *32*). The vast majority (96% in Arabidopsis and 80.1% in maize) had at least one transcription factor binding site (**Fig. 1C, fig. S2B; full statistics report in table S3, S4**). Similar enrichment for transcription factor binding and histone marks was observed in tomato (**table S4**). Only 1.065% and 0.98% of CNSs overlapped with the typical gene-body marks in plants, H3K3me1and H3K36me3, respectively (*33*). Moreover, Conservatory CNSs were highly enriched in maize super-enhancers that produce bi-directional enhancer RNAs (eRNAs) (*45*). Only 0.67% of Conservatory CNSs were unsupported by functional genomics evidence in Arabidopsis (e.g. open chromatin, transcription factor binding assay). In maize, 22.7% of CNSs lacked supporting biochemical evidence, likely resulting from less experimental data and ongoing genome re-annotations in this species. Consistent with the evolving annotation for this species, 10.2% of Poaceae CNSs detected using maize as a reference could be mapped to transposons annotated since our initial analyses (*23*).

Conservatory CNSs were significantly enriched in accessible chromatin regions in Arabidopsis, tomato, and maize (*p* < 1e-4, permutation test; **table S5**). Moreover, CNSs were preferentially located at the centers of open chromatin regions, likely pinpointing specific functional sequences within cis-regulatory space (**Fig. 1D-F**). Thus, Conservatory detects CNSs with the hallmarks of functional cis*-*regulatory elements.

We estimated the ages of CNSs by mapping them to a phylogenetic tree with nodes dated according to published best estimates. Most (2.15 million; 92%) were conserved solely within plant families. We identified many ancient CNSs that emerged prior to the diversification of the eudicots (n=31,314; 1.34%), angiosperms (n=3,321; 0.14%), or even seed plants (n=633; 0.02%) (**Fig. 1G; table S6**). We use ‘ancient’ to refer to CNSs that emerged prior to the divergence or diversification of major angiosperm clades (∼115-100 Mya) (*34*). To assess the functional categories of genes harboring the most ancient seed plant CNSs, we performed gene ontology (GO) term analysis. Similar to ancient metazoan CNSs, genes associated with angiosperm and seed plant CNSs were significantly enriched for developmental and transcriptional biological processes (*15*) (**Fig. 1H**), suggesting these ancient CNSs have critical roles in conserved developmental programs.

### Mutations in deeply conserved CNSs disrupt development

To determine the functional roles of ancient Conservatory CNS-gene associations in development, we first focused on genes with canonical roles in embryogenesis (*35*). The ancient homeobox gene *WUSCHEL HOMEOBOX 9* (*WOX9*) has conserved functions in development (*9*). We showed in tomato that CRISPR-Cas9 deletion alleles of *SlWOX9* spanning ∼600 bp of the proximal promoter cause partially penetrant embryonic lethality along with vegetative defects, including extra cotyledons, premature shoot apical meristem (SAM) termination, and stem or leaf fusions (*9*). Re-evaluation of these alleles (*9*) revealed that they disrupted an ancient seed-plant level CNS (S217). To validate these observations, we generated two additional small inversion and deletion alleles disrupting this ancient CNS, which replicated these phenotypes (**Fig. 2A-C; table S7**).

**Fig. 2.**
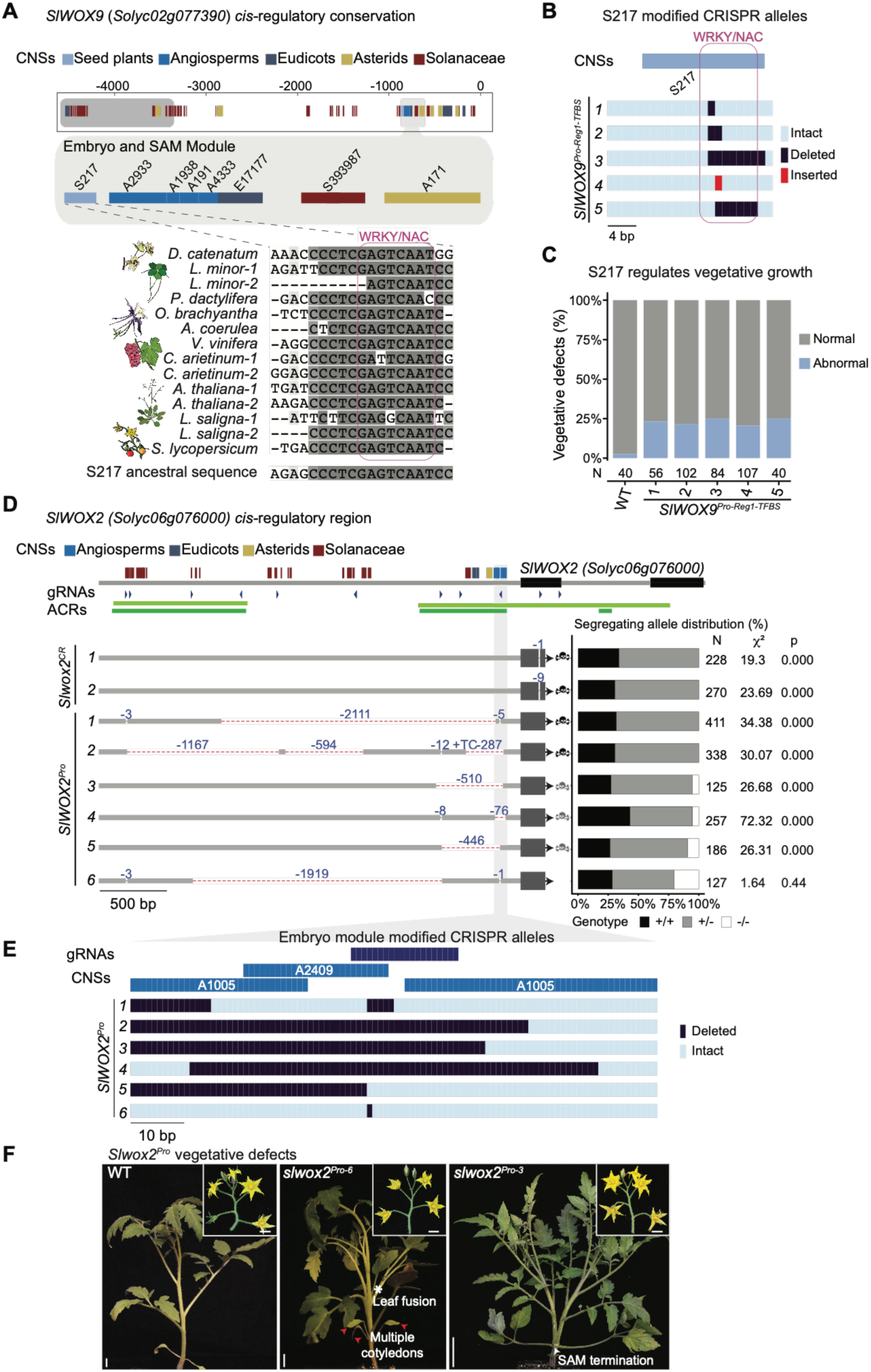
CRISPR-Cas9 mutagenesis of ancient CNSs near *HOMEOBOX* genes disrupts embryonic and vegetative development. (**A**) CNS annotations of the *SlWOX9* promoter. Distinct modules controlling expression in the embryo and shoot apical meristem (pro-Reg1) or inflorescences (pro-Reg2) (*9*) are outlined in gray. Multiple sequence alignment of seed plant CNS (S217) within pro-Reg1 for ten representative angiosperm species, with a conserved WRKY/NAC motif in pink. (See **table S7** for gene names). (**B**) Schematic of five *slwox9^Pro-Reg1-TFBS^* alleles targeting CNS S217. (**C**) Percentages of offspring with vegetative defects in wild type (WT) and in five F2 families segregating *slwox9^Pro-Reg1-TFBS^* alleles. (**D**) CNS annotations of the *SlWOX2* promoter and a *slwox2^Pro^* allelic series generated by CRISPR-Cas9. Dashed red lines indicate deletions. Black and gray skulls mark fully or partially expressive embryonic lethal alleles, respectively. The right panel shows genotypic proportions of F2 populations and chi-square statistics under a 1:2:1 wild type:heterozygous:homozygous expectation. Note the top 2 rows are *Slwox2* CRISPR null alleles. (**E**) Detailed view of deeply conserved *SlWOX2^Pro^* region, showing deletions in the different *Slwox2Pro* alleles. (**F**) Representative vegetative phenotypes of *slwox2^Pro-6^* homozygotes, with multiple cotyledons (red arrowheads) and leaf fusion (white asterisk), and *slwox2^Pro-3^* homozygotes with a terminated SAM (white arrowhead).

To expand functional analyses of ancient CNSs, we next analyzed *WOX2,* which is expressed in the Arabidopsis zygote and also acts in embryogenesis alongside *WOX9* and other *WOX* genes (*35*). To determine if *WOX2* function is conserved in tomato, we generated two independent coding null alleles in *SlWOX2*. Neither allele could be recovered as homozygotic, indicating fully penetrant embryonic lethality. This contrasts with Arabidopsis, where *wox2* mutations have no phenotypes due to redundancy with several paralogs (*35*). These divergent redundancies likely reflect the distant relationship between tomato and Arabidopsis. Conservatory identified three eudicot-level and three angiosperm-level CNSs in the *SlWOX2* promoter (**Fig. 2D**). Using CRISPR-Cas9, we generated six *SlWOX2* deletion alleles (*SlWOX2^Pro^*) of various lengths across ∼3 kb of the promoter. Five of these alleles removed all or part of the angiosperm CNSs, and all resulted in complete or near complete embryonic lethality. Similar to the vegetative phenotypes of the *SlWOX9* alleles that disrupted the S217 seed-plant CNS, rare homozygotes of the *SlWOX2* promoter alleles displayed vegetative defects. The *SlWOX2^Pro-6^* allele, where the angiosperm-level CNSs remained largely intact, showed no embryonic lethality, supporting the critical functional role of this ancient CNS (**Fig. 2B-F**) (*9*). Though gRNA availability and efficiency precluded more precise CNS perturbation, an inevitable challenge in testing CNS functionality, including associated expression changes, these alleles demonstrate key roles for one or more of these *SlWOX2* ancient CNSs in embryogenesis. The partially penetrant vegetative phenotypes suggest additional CREs contribute to these developmental programs.

Finally, we assessed the functional importance of ancient CNSs, by analyzing previously characterized CREs of eight transcription factors that regulate meristem, leaf, flower, or epidermal development (*13*, *36–41*). In all cases, functionally characterized CREs overlapped with ancient CNSs (**fig. S3**). In particular, the founding *WOX* gene, *WUSCHEL (WUS)*, a conserved regulator of stem cell proliferation (*42*), has two defined regulatory modules, previously defined by spatiotemporal expression patterns induced by promoter fragment constructs. One module drives *WUS* expression in ovules, and the other in meristems (**Fig. 3A; table S7**) (*43*). We discovered five ancient CNSs in the meristem module, including conserved binding sites for Type-B RESPONSE REGULATORS (RRs), which act downstream of cytokinin to induce *WUS* expression (**Fig. 3A, fig. S4A; table S7**) (*44*, *45*). This strong correspondence between classical developmental CRE studies and Conservatory CNSs further supports the importance of these sequences.

**Fig. 3.**
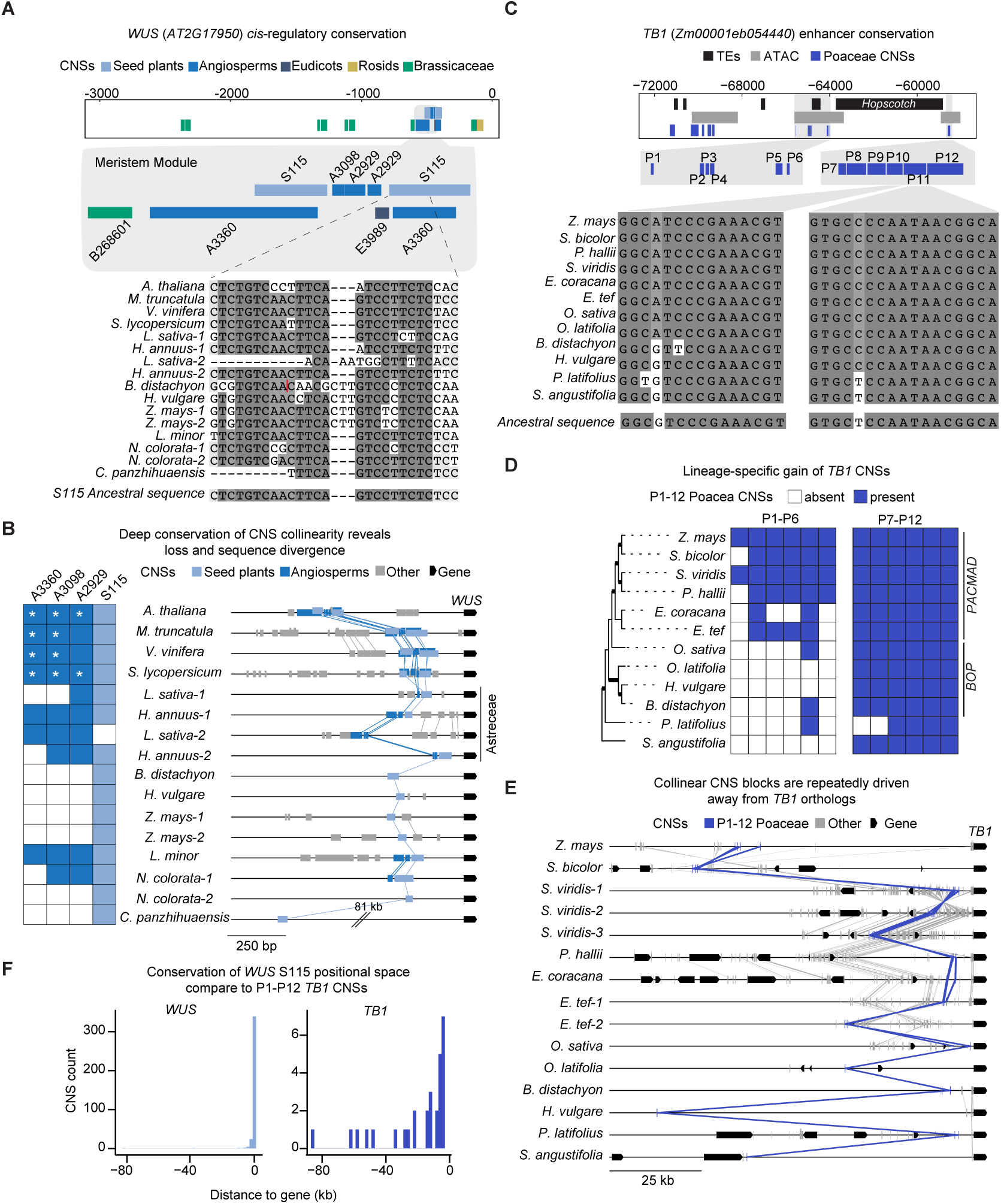
CNSs associated with homologs of *WUSCHEL* and *TEOSINTE BRANCHED1* illustrate principles of plant *cis-regulatory* evolution. (**A**) CNS annotation of the *AtWUS* promoter. The conserved regulatory module specifying *WUS* expression in the SAM (*43*) is shown at high detail, with multiple sequence alignment of the seed plant CNS S115 (partial) for twelve representative angiosperm species (see also **fig. S4**). The red line marks the position of a *B. distachyon-*specific insertion that was omitted for visualization purposes. (**B**) Presence-absence matrix (left) and microsynteny plot (right) of deeply conserved CNSs in the *WUS* meristem module. Asterisks indicate intragenic duplication of a CNS. (**C**) CNS annotation of the *ZmTB1* promoter. ACRs on either side of a *Hopscotch* transposon each contain six CNSs conserved in the grass family. Shown below are multiple sequence alignments of CNSs P10 and P11 in twelve selected grass species, and the presence-absence matrix of these CNSs in (**D**). P1-P6 are specific to one clade, while P7-P12 are broadly conserved among grasses. (**E**) Microsynteny plot of *TB1* orthologs. The P1-12 block is highlighted in purple. Note the varied position of this CNS group in different grasses. (See **table S7** for gene names, and **table S8** for P1-P12 block coordinates). (**F**) Distributions of CNS distances from the translation start sites of *WUS* and *TB1* orthologs.

### Two core meristem regulators uncover CNS evolutionary dynamics

We used Conservatory’s identification of CNS homologies to investigate how CNSs evolve, first focusing on the deeply conserved *WUS* meristem regulatory module. In Arabidopsis, Conservatory identified five partly overlapping CNSs in this module. The S115 CNS sequence, deeply conserved even in the gymnosperm *Cycas panzhiuaensis*, was duplicated and identified in distal and proximal positions (**Fig. 3A; table S7**). Cross-species analysis suggested that the core spatial configuration of this meristem module emerged at the base of the angiosperms, while the S115 duplication likely occurred at the base of the eudicots. While 3 of the 4 CNSs in this module were lost in the grasses, the deeply conserved S115, which contained a Type-B RR binding site important for meristem expression (*44*, *45*), was retained (**Fig. 3B; table S7**). Consistently, the *WUS* meristem expression pattern is conserved in the grasses, despite divergent meristem morphology and developmental dynamics (*46*, *47*).

*WUS* underwent several independent duplications, and CNS analysis for the duplication that occurred at the base of the Asteraceae showed that associated cis*-*regulatory sequences were duplicated with the coding regions. However, these sequences then exhibited species-specific patterns of divergence and loss (**Fig. 3B**). In lettuce (*Lactuca sativa)*, only one angiosperm-level CNS was retained in both *WUS* paralogs, while the other ancient CNSs were differentially retained and partitioned between the two paralogs. The products of the same ancestral duplication diverged differently in sunflower (*Helianthus annuus)*, with one *WUS* paralog maintaining all ancient CNSs, and the other maintaining 3 out of 4 of them (**Fig. 3B**). Both species appeared to have lost the distal copy of S115, but closer examination revealed an Asteraceae-specific CNS similar to the ancestral S115 sequence in the collinear location (**fig. S5A-C; table S7**), suggesting sequence divergence followed by sequence maintenance in the Asteraceae. Analysis of an additional *WUS* duplication that occurred at the base of a large clade of grasses revealed similar evolutionary patterns, with S115 and Poaceae-specific CNSs shared between the paralogs, but each paralog also had lineage-specific CNSs (**fig. S5D; table S7**). Despite divergence in the regulatory sequences of *WUS* paralogs, the ancient meristem module CNSs were maintained across angiosperms (**Fig. 3B**).

To characterize CNS evolution over shorter time scales, we next analyzed the maize *TEOSINTE BRANCHED1* (*TB1*) gene. *TB1* encodes a TCP transcription factor with conserved roles in axillary branching (*48–50*). Regions upstream of *TB1* orthologs are associated with maize (*Zea mays*), sorghum (*Sorghum bicolor*), and pearl millet (*Pennisetum glaucum*) domestication (*51–53*). Analysis of the maize domestication locus, located ∼65 kb upstream of *TB1,* uncovered multiple CNSs, separated into distal and proximal collinear clusters (P1-P6 and P7-P12, respectively; **Fig. 3C; table S7 & S8**). Tracing the evolution of *TB1* CNSs revealed that the proximal cluster emerged in a lineage leading to the grasses (∼70 mya), while the distal cluster emerged later, before the radiation of a major grass clade that includes maize (PACMAD clade, ∼33 mya, **Fig. 3D**). A gain-of-function *Hopscotch* transposon insertion associated with domestication separates the proximal and distal CNS clusters in maize, leaving the CNSs intact. Individual fragments of the regulatory region containing these CNS act as repressors (*54*), suggesting that the gain-of-function *TB1* phenotype in maize could be caused by the *Hopscotch* insertion disrupting CNS spacing in a conserved repressor element (**Fig. 3A, E**).

Comparison of CNS spacing and order in *TB1* and *WUS* CNSs revealed that local CNS collinearity was retained across species, with the exception of an inversion in several grasses (**Fig. 3E**). In contrast, the distances between CNSs and their associated genes varied and were CNS-specific. Thus, the *WUS* meristem CNS module was within 1.5 kb of the start codon in 224 of 240 species, and only in 8 cases was more than 10 kb upstream. The position of the *TB1* enhancer was much more variable (**Fig. 3F**), and in some cases, such as sorghum, it was closer to a different gene than to *TB1* itself. In other grasses, such as *Oryza latifolia*, a different gene was located between the CNS cluster and the *TB1* ortholog coding region (**Fig. 3E**).

Overall, our analyses of *WUS* and *TB1* revealed that CNSs maintained after duplication diverge in particular lineages, change position relative to their associated genes, or even get lost as cohorts in specific lineages. Gene duplication was associated with duplication of associated CNSs, which remained as multiple copies in the genome. We next asked whether such dynamics characterized CNS evolution on a global scale.

### Genome rearrangements alter CNS position and gene associations

To study the evolutionary dynamics of CNS position, we first analyzed the positions of all CNSs relative to their associated genes, grouped by CNS age. The distribution of CNS positions was genome-specific, but overall, 73% of CNSs were located upstream of associated genes. They were enriched near coding sequences, with angiosperms and seed plant-level CNSs median position being closer to the start codon than family-level ones (2,252 bp vs 11,865 bp, respectively; **Fig. 4A**; *p* < 0.001; Wilcoxon rank sum test). However, 25% of CNSs across all age categories were more than 25 kb from their associated genes. In 47.5% of cases, CNSs were located closer to a different gene than to their associated gene. While the precise number was genome specific, between 25% to 50% of CNSs-gene associations skipped over intervening, non-associated genes (**Fig. 4B; table S9**).

**Fig. 4.**
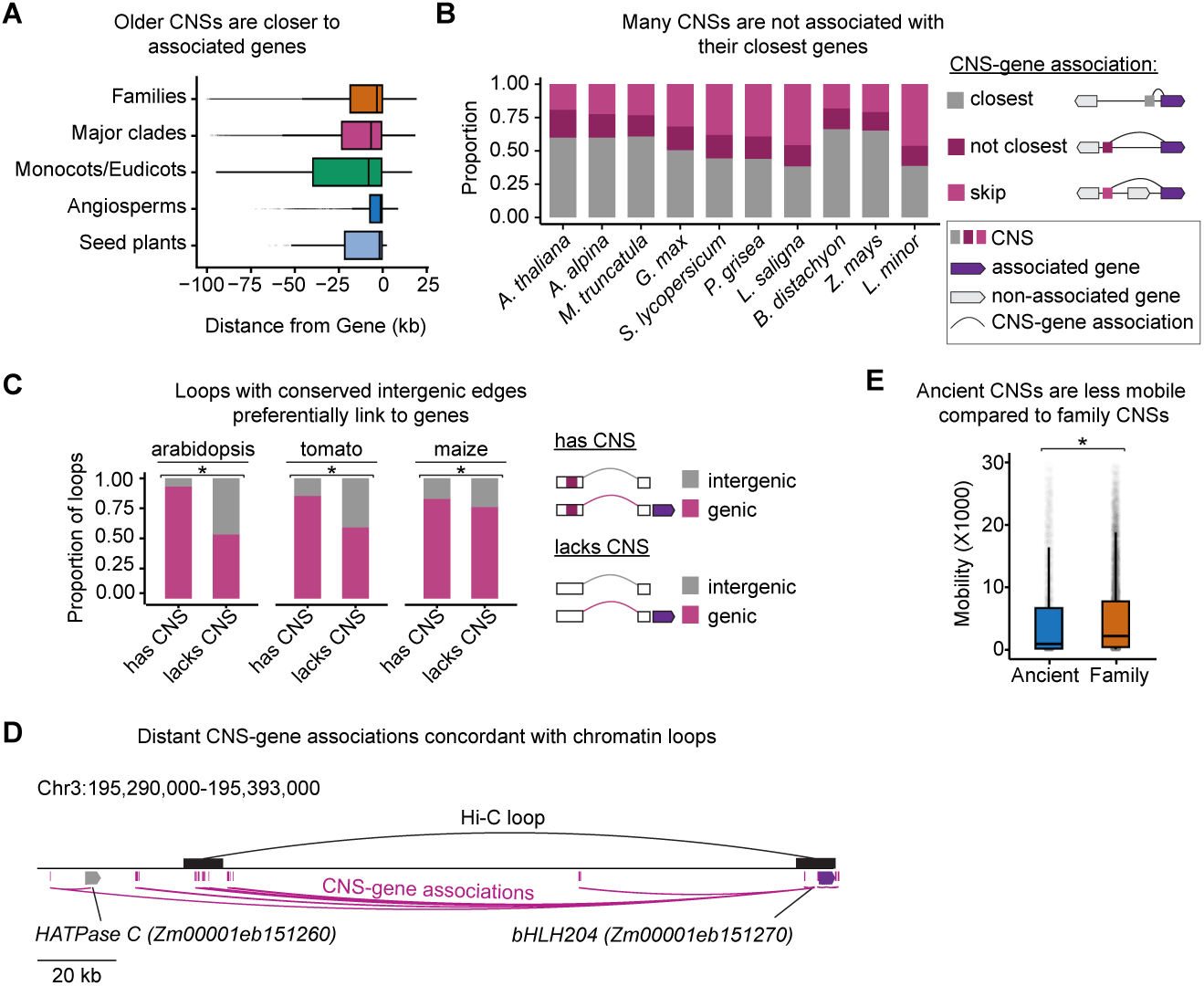
Conservation of CNS position and gene associations. (**A**) Distributions of CNS-gene distances, stratified by CNS conservation level. (**B**) Proportions of CNSs associated with the closest genes, with other adjacent genes, or that skip adjacent genes, for each of our ten reference genomes (**table S9**). (**C**) Proportions of intergenic chromatin loop edges in Arabidopsis, tomato, and maize, stratified by overlap with at least one CNS and whether the other edge of the loop connects to a gene. Chromatin loops connecting CNSs with genes are more prevalent than those that do not overlap CNSs (*p <* 0.001; chi-square test, **table S10**). (**D**) Example of long-range CNS-gene association supported by chromatin looping. (**E**) Distribution of CNS mobility (standard deviation of relative position in orthogroups of taxonomic family) of ancient (angiosperm and seed plant) vs family level CNS (p<2.2e-16; Wilcoxon test).

Long-range CNS-gene interactions and interactions skipping over other genes are common in metazoans (*55*), but are only rarely reported in plants. While they may be common in the large maize genome (*56*), we found many distal or non-adjacent associations, including in compact genomes. In Arabidopsis, tomato, and maize, 74-88% of genes intervening between CNSs and their associated genes were associated with CNSs of their own, and for 95-96% of gene-skipping CNSs in these species, the closest non-associated genes were not orthologous to associated genes. This suggests that CNS-gene associations that skip over non-associated genes were unlikely to be caused by annotation artifacts or by tandem gene duplications (**fig. S6A-B**). To provide empirical support for such long-range CNS-gene associations, we investigated the concordance of predicted CNS-gene associations with predicted chromatin loops, as cis-regulatory elements and gene interactions might be maintained through chromatin looping (*56*). We carried out chromatin-conformation capture sequencing on tomato meristem tissue, and reanalyzed published datasets for Arabidopsis and maize, focusing on loops connecting intergenic regions to genes (*56–58*). Compared to loops that did not overlap with CNSs, loops with CNSs at their intergenic edges were more likely to be connected to genes (*p* < 0.001, chi-square test) (**Fig. 4C-D; table S10**). This suggests that intergenic DNA sequences looping to genes are conserved, consistent with Conservatory CNSs acting as regulatory elements. Although conserved synteny does not necessarily imply regulatory function, our analyses support the long-term persistence of synteny between genes and their associated CNSs.

Our analysis uncovered an unexpected relationship between the depth of CNS conservation and relative position to their associated genes, with ancient CNSs occurring closer to transcription start sites. One explanation for this observation is that the functions of these CNSs are position-dependent. In support of this, analysis of CNS mobility (defined as the standard variation of its relative position) over short evolutionary distances (i.e., within taxonomic families) revealed that mobility of ancient CNSs was still significantly lower than that of comparable family-level CNSs (p<2.2e-16; Wilcoxon test; **Fig. 4E**).

### Evolution of new cis*-*regulatory sequences from ancestral CNSs

Our analysis of the *WUS* meristem module revealed that even ancient CNSs can diverge or be lost in specific lineages. To assess the extent of this phenomenon, we examined the presence or absence of ancient CNSs across species. The number of angiosperm-level CNSs detected in each species varied widely, yet was highly consistent within families, suggesting that most losses or divergences occurred during early events at the base of each family (**Fig. 5A**). CNS absence in individual genomes may result from incomplete genome assembly or misannotation, but the consistency of CNS presence across independent species within families suggests that these CNS loss events are a biological phenomenon, rather than technical artifacts. Using a maximum parsimony approach, we mapped the most likely phylogenetic nodes where angiosperm CNS loss or divergence occurred. While there was a gradual overall trend of CNS loss or divergence, we also detected punctuated, large-scale losses in families such as Brassicaceae, Asteraceae, Poaceae, and Lemnaceae (**Fig. 5B**). In the grasses, this large-scale CNS loss is consistent with rapid evolutionary diversification (*59*). Indeed, lost CNSs were associated with the *WOX4* and *KANADI* transcription factor genes, which play key roles in vasculature and leaf development, processes in which grasses display unique morphological innovations (**table S11**) (*60*).

**Fig. 5.**
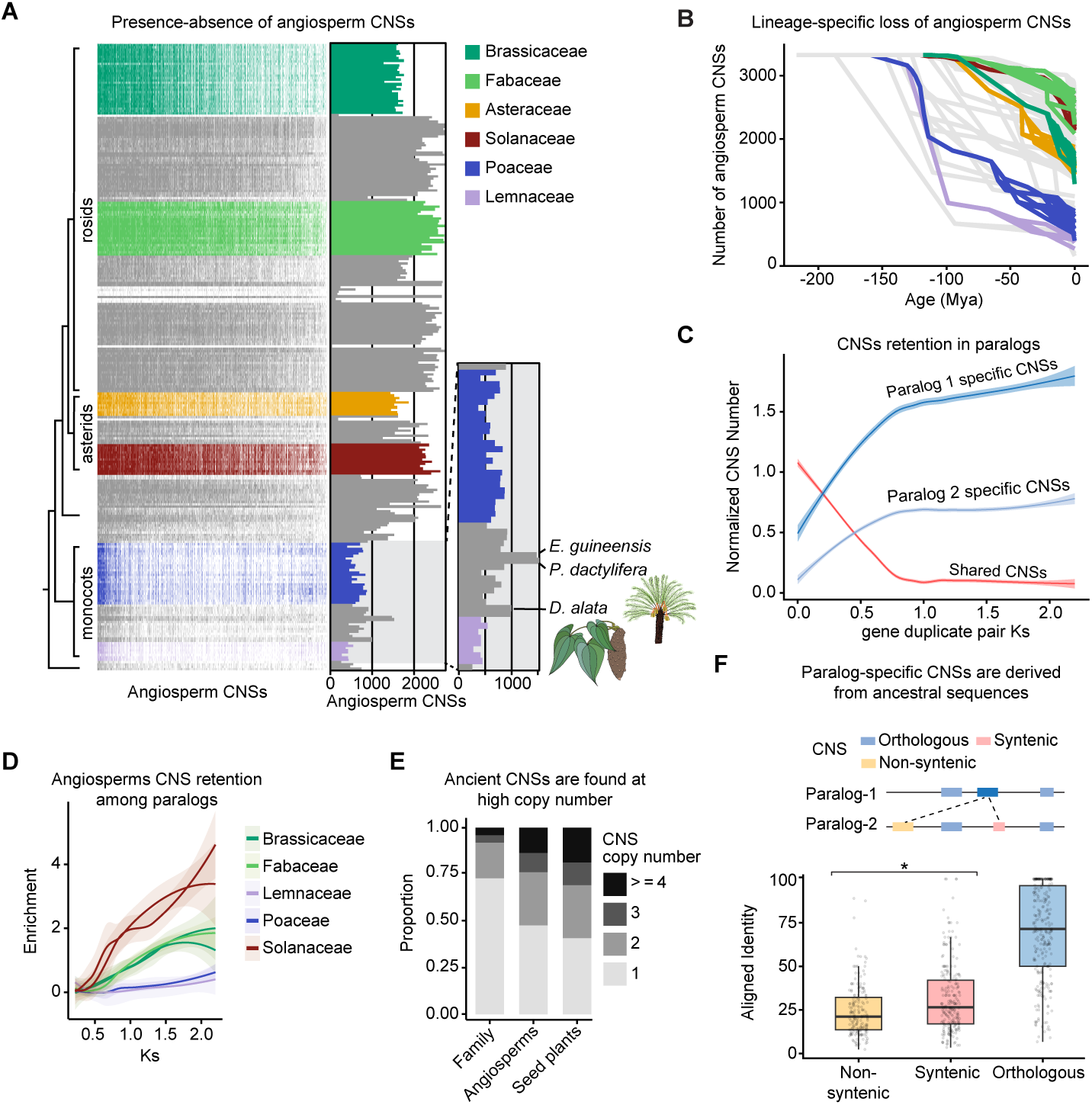
Diversification of CNSs throughout plant evolution. (**A**) Presence-absence matrix of angiosperm CNSs among all species in this study. Rows correspond to genomes, columns to angiosperm CNSs, and tiles are color-coded by taxonomic family. The lower right panel highlights outlier monocot species. The boxed region is a histogram summarizing the presence-absence calls. (**B**) Lineage-specific loss of angiosperm CNSs. Each line tracks the number of angiosperm CNSs in a single genome from 200 Mya to the present day. Lines are color-coded according to the taxonomic family. (**C**) The normalized CNS number in diverging paralogs resulting from whole-genome duplications in 10 reference genomes. Shown is the number of shared CNS amongst the paralogs, paralog-specific CNS, and the number in each of the paralogs. To control for variation in CNS number between genomes, the CNS count per gene was normalized to the mean CNS count per gene in the relevant genome. (**D**) Enrichment of angiosperm-level CNS retention in duplicate gene pairs from the reference genomes relative to simulated random CNS retention. Two genomes were used from each family. As paralogs diverged, the preference to maintain the angiosperm-level CNS increased. (**E**) Distribution of CNS copy number in genomes stratified by CNS conservation level. (**F**) Distribution of pairwise identity for nonsyntenic CNSs, for syntenic CNSs with different names, and syntenic orthologous CNSs. Syntenic CNS were more similar than non-syntenic ones (p<0.001; Wilcoxon test).

One explanation for the apparent loss of ancient CNSs may be CNS divergence accompanied by the emergence of lineage-specific CNSs, as for *WUS* in the Asteraceae (**fig. S5**). To assess CNS divergence more broadly, we analyzed paralog evolution within genomes as a proxy for early stages of gene evolution. We focused on paralog evolution because sparse phylogenetic sampling limits detection of early divergence events across species. Analyzing paralogs allows ancestral CNS complements to be reconstructed and regulatory divergence directly traced. We identified duplicate pairs derived from whole-genome duplications in our 10 reference genomes. Following duplication, the number of shared CNSs declined with age, as the number of paralog-specific CNS steadily accumulated. The acquisition of paralog-specific CNS was asymmetric, with one paralog being dominant. As coding sequence diverged CNS number reached a plateau, revealing an early phase of divergence marked by CNS loss and divergence that was subsequently stabilized (**Fig. 5C**). As for *WUS* paralogs, ancient CNSs were preferentially maintained in both paralogs compared to younger, family-level ones. Consistently, these old CNSs accumulated in higher copy numbers in genomes (**Fig. 5D-E**).

Finally, we asked whether new paralog-specific CNSs arise de novo or are derived from ancestral sequences (*61*). Distinguishing de novo CNS formation from preferential retention is challenging without dense phylogenomic coverage. We therefore focused on paralog pairs in which an old CNS (present before duplication) was retained in one paralog but replaced by a different CNS at the same position in the other paralog (**Fig. 5F**). We reasoned that if these sequences are derived, they should still retain some sequence similarity to the old CNS. Using CNSs retained in both paralogs as alignment anchors, we compared the sequence similarity between old CNSs and new paralog-specific CNSs at syntenic sites versus those in non-syntenic sites. Syntenic replaced–new CNS pairs showed significantly higher identity than non-syntenic controls (**Fig. 5F**; p < 0.001; Dunn test). Thus, although prevalence remains to be assessed, our analysis suggests that new paralog-specific CNSs can originate from ancestral sequences rather than arising de novo.

## Discussion

Conservation and divergence of cis-regulatory sequences are key forces driving morphological evolution, yet their evolutionary dynamics have remained poorly understood. Using Conservatory as a platform for CNS discovery, we identified and traced the evolution of over 2 million CNSs in plants, including many that predate the emergence of angiosperms.

Why are these sequences so deeply conserved? Their enrichment near developmental regulators suggests they govern critical regulatory programs. This is supported by the severe developmental phenotypes observed when such sequences are disrupted (**Fig. 3**) (*9–12*, *14*). Moreover, their retention in high copy numbers following gene duplication suggests that their function may be less dependent on the surrounding sequence environment. Thus, similar to ancient metazoan CNSs, ancient plant CNSs may encode functionally-enriched core regulatory sequences (*62*). The deep conservation of these sequences does not imply that these are the major or only regulatory elements. Rather, the observed gradient of CRE selective constraints, with slowly evolving deeply conserved sequences flanked by faster evolving sequences, suggests a model whereby deeply conserved sequences are a bedrock of complex cis-regulatory control of expression programs, which are tuned by more shallowly conserved peripheral sequences, whose modifications contribute to trait variation and innovation.

Following gene duplication, the regulatory landscapes of the resulting paralogs rapidly diverge (*63*, *64*). Analysis of the specific *WUS* and *TB1* duplications reveal lineage and gene-specific divergence patterns, but overall, there is a clear asymmetry between the paralogs, with one being dominant in terms of CNS number. This may represent one paralog contributing to gene family diversification through sub- or neo-functionalization, but even diverged paralogs tend to maintain some shared CNS space, suggesting retention of partial redundant function.

Although new regulatory sequences are often thought to arise de novo from neutrally evolving noncoding DNA, many may originate from older sequences (*5*, *15*, *61*, *65*). By tracing CNS dynamics in paralogs, we provide direct evidence that new CNSs often emerge through divergence of ancestral elements rather than de novo formation. Furthermore, this suggests that certain sequences, and the composition of TFBSs within them, possess intrinsic properties that predispose them to regulatory function and long-term retention.

One of the paradoxes in CNS evolution is that deeply conserved sequences can nevertheless be lost in cohorts within specific lineages. Such loss may result from extreme developmental systems drift, in which, after gene duplication or divergence of other regulatory sequences, a deeply conserved CNS becomes redundant and gradually degenerates in function or becomes a derived CNS through mutation (*66*, *67*). Alternatively, it may reflect technical limitations: orthologous CREs are difficult to align and identify across large evolutionary distances (*17*, *18*). While leveraging hundreds of genomes in this study helped bridge these gaps, sparse phylogenomic coverage remains a constraint for reliably identifying CNS orthology. For example, the apparent loss of many deeply conserved CNSs in grasses and duckweeds is consistent with large-scale cis*-*regulatory rewiring during their early divergence (*23*, *68*, *69*). Yet poor sampling of monocots outside these families means that many lineage-specific CNSs may still be orthologous to ancestral angiosperm elements. Our study provides a unified map of plant CNS evolution based on 314 genomes. As pan-genomic resources expand, improved phylogenomic resolution will further clarify orthologous relationships between CNSs and guide deeper computational analyses and functional assays to refine our understanding of regulatory sequence evolution and provide even higher resolution targets for trait engineering (*5*, *70*).

## Materials and Methods

### Conservatory algorithm

The Conservatory algorithm was written in perl and R. All scripts, installation instructions and related data files are available at https://github.com/idanefroni/Conservatory/. All references to scripts below are to those in the github scripts/ directory. Output files, including raw data, calibrated trees, and GFF3 tracks for all 314 genomes used in this study, are provided at www.conservatorycns.com. The conservatory algorithm was run on the Hebrew University High Performance Computing cluster, using up to 1,000 nodes at a time. Overall run time for producing and merging the entire dataset was approximately 7 days. Jobs used to run the algorithm are provided in the jobs folder of the GitHub directory. Parameters for alignment sensitivity, filtering thresholds, and search-space length were empirically determined based on their ability to identify a curated subset of known cis-regulatory elements (CREs) and are provided in **table S12**.

### 1. Genome resources

Genome assemblies and annotations were downloaded from public repositories, including NCBI, Ensembl, CoGe, Phytozome, and project-specific websites as described in **table S1**. We analyzed 314 genomes from 68 plant families. For each genome, we obtained FASTA format files containing genome and protein sequences, and GFF3 gene annotation files. Ten genome assemblies were designated as family-specific reference genomes: *Arabidopsis thaliana* and *Arabis alpina* for Brassicaceae, *Medicago truncatula* and *Glycine max* for Fabaceae, *Lemna minor* strain 7210 for Lemnaceae, *Brachypodium distachyon* and *Zea mays* for Poaceae, *Lactuca saligna* for the Asteraceae, and *Solanum lycopersicum* and *Physalis grisea* for the Solanaceae.

For these ten reference genomes, we generated repeat annotation using EDTA (*71*) with default settings. BUSCO completeness of each genome was assessed using BUSCO v6.0.0 (*72*). For each genome, the assembly and predicted proteins were searched against both embryophyta_odb12 and viridiplantae_odb12 reference datasets with miniprot v.0.18-r281 (*73*) and HMMER v3.4 (*74*). Genome assembly statistics were calculated within the BUSCO pipeline using BBTools version v.39.33 (*75*) (**table S1**).

### 2. Phylogenetic inference

Conservatory requires two types of phylogenetic trees. First, family-specific rate-calibrated trees are required for each reference species. Second, a single all-species tree with age estimation for each node. To generate phylogenies for each of the reference families, we used our initial orthogroup assignments to identify low and single-copy nuclear genes in each family. The predicted peptide sequences for each of these orthogroups were aligned with mafft (v7.313) (*76*). Alignments were trimmed with clipkit (*77*) and concatenated. Because the taxonomic sampling of available genomes was inappropriate for accurate phylogenetic inference, and because our primary goal was not to determine relationships between species, but to use best estimates from systematics, constraint trees were generated for all families using published phylogenies (*78–106*). Maximum likelihood tree inference was then carried out with RAxML-NG version 1.1.0 (*107*), providing the concatenated orthogroup alignments, a constraint tree, and partitions, and specifying a JTT+G substitution model.

To generate the all-species tree, we clustered the predicted peptide sequences from each genome into orthogroups using OrthoFinder (*108*). To identify low-copy orthogroups suitable for species tree construction, we aligned tomato peptide sequences from the unmasked multiple sequence alignments of the 1KP project (*109*), to tomato SL4.0 predicted peptides with blastp at default settings, retaining the top-scoring hit for each 1KP peptide. This recovered 386 of the 410 1KP orthogroups. As a quality control, we inspected the distribution of these orthogroups present in each genome, and the number of genomes with each orthogroup. Among 160 orthogroups present in 300-500 total copies among all genomes, we selected 84 orthogroups at random.

Another 16 orthogroups were specifically chosen for *Apium graveolens* and *Nelumbo nucifera* genes, for a total of 100 orthogroups. Alignments and phylogenetic inference were conducted as for the family trees, with relationships between major clades constrained following (*109*), except phylogenetic inference was performed using IQTree2 version 2.0.3 (*110*). Node ages were estimated with the penalized likelihood approach (*111*, *112*) implemented in the R package ape (*113*). We used a relaxed model of substitution rate variation among branches and up to 80 alternative iterations between estimating rates and dates. Calibration points were obtained from TimeTree (*114*) and from the literature (*78–106*) and are available as **table S14**.

### 3. Genome and mapping database construction

Whole-genome sequences and gene annotations (in GFF3 format) were preprocessed using the <u>processGenomes</u> script, which standardized FASTA headers, indexed sequences for rapid retrieval, and parsed gene names to generate standardized gene coordinate files. Repetitive sequences for all reference species were masked based on the EDTA annotation (see above). Regulatory sequences were extracted using the <u>buildRegulatorySeqDB</u> script. For each gene, non-coding regions were extracted from the genome. The length of upstream regulatory regions was determined as the 97th quantile of intergenic region lengths, with a maximum of 100 kb. The downstream regulatory region was one-fifth the length of the upstream region. Upstream sequences were taken from the translation start site, and downstream from the translation end site. To prevent contamination by coding sequence, CDSs were masked based on genome annotation.

Homolog databases were created using the <u>buildOrthologDB</u> script. First, reciprocal all-vs-all BLAST against all other genomes in the database was performed. Each gene pair was accepted as orthologous if each gene was the top BLAST hit of the other in the corresponding search direction. In cases where multiple high-scoring hits existed, a cut-off E-value was calculated at 0.66 of the maximum E-value score. Up to 16 genes with an E-value higher than the cutoff were defined as a group of homologs. Sequences shorter than 120 amino acids were treated with higher sensitivity, using relaxed E-value cutoffs. The final output was a tab-delimited table containing homolog groups with associated alignment scores.

### CNS identification

Identification of CNS for a given gene was performed using <u>buildConservation</u> script. For each gene in the reference species, candidate homolog genes were identified in the target species using the homolog mapping database. The regulatory regions of the gene were filtered for low complexity regions and then aligned using LastZ (--format=maf --gap=200,100 --nochain --noytrim --seed=match4 --gapped --strand=both --step=1 --ambiguous=iupac --identity=70 --ydrop=1000 --hspthreshold=2000 --gappedthresh=2000) and alignment scores calculated. The highest-scoring homolog was selected as the putative ortholog; however, if multiple candidates had comparable high scores, multiple orthologs were retained per species to account for recent duplications or lineage-specific expansions. To increase the usability of genomes with incomplete assembly or annotation, multiple genomes can be assigned for one species. In this case, Conservatory selects species that had multiple available genomes, with the homolog from the genome with the best alignment score for the regulatory regions selected for further analyses.

Alignments were refined by trimming poorly aligned ends and merging nearby high-scoring alignment blocks. Conservation within the aligned sequences was assessed using *phyloP* (*27*), using the family tree, with rates estimated from the background substitution rates, as detailed above. Conserved blocks were defined as regions with phyloP p-values lower than a species-specific p-value cut-off and over stretches of at least 15 bp. The species-specific values were selected so that, on average, the number of conserved bp per gene is 250. Candidate conserved blocks were filtered for a minimum sequence identity of over 70% and a minimum length threshold of 8, and regions overlapping masked or low-quality genomic regions were removed.

For each CNS, ancestral sequences were reconstructed. First, sequences were aligned using mafft (--auto --thread -1 --ep 0.3), followed by *FastML* (-b -qf -mh) using the family tree. Then, each reconstructed ancestral sequence was aligned with homologous gene regulatory sequences of all genomes from outside the family using sensitive local alignment with relaxed parameters (--HSPThreshold=1600). Hits were retained if they met identity (> 70%) and length (> 12 bp) thresholds.

To obtain a unified CNS map for a reference genome, the <u>mergeCNS</u> script was used. CNS entries were compared based on genomic coordinates, and overlapping CNSs were collapsed into a single entry, prioritizing longer sequences, higher conservation scores, and broader conservation across species. A final filtering step was used to remove low-quality and spurious CNS associations.

Following the merge of the CNSs, sequences that result from unannotated coding sequences were filtered. CNS sequences were used for a tblastn against a combined protein dataset of all annotated proteins in the 314 genomes. CNSs that had more than 5 hits at Evalue lower than 0.0001, or had longer than 200 aligned basepair were considered as unannotated open reading frames and were removed from the dataset using the scripts <u>blast_filter.sh</u>.

To obtain a single CNS map for multiple reference species, the CNS dataset files were concatenated and then merged using the <u>mergeCNS</u> script with the --merge-deep parameter. This algorithm searches for overlapping CNSs from different reference species. If a sufficient number of CNSs overlap, they are merged to form a single CNS. The combined sequences are then realigned using mafft and then split or merged based on the depth of conservation. The CNS-gene association is filtered based on phylogenetic relationships. Finally, ancestral sequences are reconstructed, as detailed above, and CNS age was determined based on the deepest node in the all-species tree common to all species with the CNS. The deepest node was defined as that which led to a clade that included all those genomes that contain a particular CNS. Here, we assumed conservation as more likely than pervasive, frequent convergent evolution of locally collinear cis-regulatory sequences, in synteny with homologous genes. CNS ‘Collinearity’ refers to local instances when Conservatory CNSs occur in the same order across multiple genomes.

### Assay for transposase-accessible chromatin using sequencing (ATAC-Seq) libraries

ATAC-seq was performed as previously described (*9*). Briefly, six biological replicates of 1g of freshly collected terminal leaflet of young M82 leaves, were ground in ice-cold isolation buffer (300 mM sucrose, 20 mM Tris pH 8.0, 5 mM MgCl_2_, 5 mM KCl, 5 mM beta-mercaptoethanol, 35% glycerol, 0.2% Triton X-100). After grinding, the suspension was filtered through a series of cell strainers (the smallest being 70 mm). Samples were further disrupted using a Dounce homogenizer, and then washed four times with ice-cold isolation buffer. The enriched nuclei samples were resuspended in Resuspension Buffer (50 mM Tris pH 8.0, 5 mM MgCl_2_, 5 mM beta-mercaptoethanol, 20% glycerol). In order to continue with 100,000 nuclei, a small portion of the nuclei-enriched sample was stained with 4,6-Diamidino-2-Phenylindole (DAPI), and counted under a microscope using a hematocytometer. For the transposase reaction 100,000 nuclei were resuspended in 22.5 ml freezing buffer (5 mM MgCl_2_, 0.1 mM EDTA, 50 mM TRIS pH 8.0, 40% glycerol), and were mixed with 2X DMF Buffer (66 mM Tris-acetate (pH 7.8), 132 K-Acetate, 20 mM Mg-Acetate, 32% DMF) and 2.5ul Transposase enzyme from Illumina Nextera Kit (Catalog No. FC-121-1031), and incubated at 37°C for 30 min. Tagmented DNA was isolated using NEB Monarch PCR Purification kit (Catalog No. NEB #T1030). Next-generation sequencing (NGS) libraries were amplified in two steps (*115*) using KAPA HiFi HotStart ReadyMix PCR kit (Catalog No. KK2601). NGS libraries were PE50 sequenced on an Illumina MiSeq SY-410-1003 instrument.

ATAC-seq reads were trimmed with trimmomatic (*116*) and aligned to Heinz SL4.0 reference genome (*117*) with BWA mem (*118*). Reads that mapped to either chloroplast or mitochondria were filtered out together with PCR duplicate reads (picard MarkDuplicates) and multiple-position aligned reads. Finally, only high-quality properly paired reads were retained for the analysis (samtools view -b -h -f 3 -F 4 -F 8 -F 256 -F 1024 -F 2048 -q 30). ATAC-seq peak calling was done by Genrich V0.6 (https://github.com/jsh58/Genrich) using all six libraries with the following parameters: -j -r -v -q 0.5 -g 30.

### Definition of background regions

To calculate CNS enrichment for functional genomic marks, we defined an appropriate background space as the genomic regions that Conservatory searches for CNSs. This is a genome-specific length upstream and downstream of all protein-coding genes, excluding annotated transposons and genes within this space. For each gene, we identified the regions spanning the start and stop codons of the longest splice isoform, then used bedtools slop to expand these regions upstream of start codons by the 97.5th percentile of intergenic distances in the genome at hand, up to a maximum distance 100 kb, and downstream of the stop codon by 20% of the upstream distance, rounded to the nearest 1 kb. Genome-specific distances are provided in **table S1**. Next, from each expanded window, we used bedtools subtract to remove regions corresponding to protein-coding genes, transposable elements and assembly gaps. The resulting gene-specific intervals were merged to delineate each genome’s background space.

### Functional genomic marks

Coordinate files in GFF3, BED, bigwig or narrowPeak format were retrieved from original publications or public databases and lifted over to different versions of genome assemblies as needed using CrossMap (*119*), as specified in **table S3**.

Maize ChIP-seq peaks (*120*) in B73 version 4 coordinates were retrieved from NCBI Gene Expression Omnibus (GEO) and converted to B73 version 5 coordinates, using the B73 v4 to v5 chain file available from Ensembl release 61. Maize DAP-seq in v5 coordinates (*121*, *122*) was obtained from GEO. Other maize epigenomic marks (histone modifications, ATAC-seq, bisulfite-seq) in B73v5 coordinates were obtained from MaizeGDB. Maize DNA methylation in bigwig-formatted coverage and methylation tracks were converted to bedgraph with bigWigToBedGraph (*123*). Transcription factor binding sites were predicted for the maize genome with PWMscan (*124*) for all nonredundant position weight matrices in the JASPAR core 2024 set, using a cutoff of 1e-5. Predicted binding sites were subsequently filtered to retain only those sites with the highest observed score for each PWM.

Arabidopsis ATAC-seq (*125*, *126*) datasets were downloaded from the Plant Chromatin State Database (*127*) as bigwig files and converted to bedgraph format for peak calling with MACS3 bdgpeakcall (*128*). Arabidopsis TF and histone ChIP-seq peaks were downloaded from the REmap 2022 database and used without additional processing (*129*). Arabidopsis DNA methylation data (*130*) was downloaded from NCBI GEO in wig format. Histone modification data were downloaded from PCSD (*127*) and broad peaks identified with MACS3 (*128*).

Published tomato meristem ATAC-seq (*9*) and histone ChIP-seq peaks (*131*) were obtained from NCBI GEO. Histone ChIP-seq peaks in M82 v1 assembly coordinates were lifted over to SL4.0 with CrossMap (*119*). Tomato TF ChIP-seq raw reads available at NCBI SRA under project ID PRJNA545571 were processed using nf-core/chipseq v2.0.0 (*132*). Briefly, raw ChIP-seq reads were trimmed using Trim Galore v0.6.10 (*133*) and aligned to SL4.0 with bwa mem v0.7.17 (*118*). Duplicates were marked with picard v2.27.4 (*134*). Alignments were filtered using samtools v1.15.1 (*135*) to remove duplicates, reads not marked as primary alignments, unmapped reads, and reads mapping to multiple locations; with bamtools v2.5.3 (*136*) to remove reads containing >4 mismatches and read pairs with insert size >2kb or in non-FR orientation. Peaks were then called from processed alignments with MACS2 (*128*). For enrichment calculations, TF ChIP-seq peaks were merged with bedtools v2.31.1 (*137*). Processed bisulfite sequencing data for tomato available from NCBI GEO under series GSE285260 were downloaded and converted to bedgraph with bigWigToBedGraph (*123*).

For each functional genomic mark, we counted how many unique CNS base pairs overlapped with that feature and with the background regions described above. Then, enrichment values were calculated as:

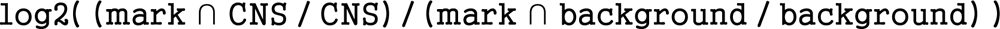

To determine the statistical significance of CNS overlaps with different functional marks, we constructed 2x2 contingency tables from the base pairs in Conservatory-sequence space or 2 kb upstream of genes that either 1) overlapped both a CNS and the mark at hand, 2) overlapped a CNS but not a mark, 3) overlapped a mark but not a CNS or 4) overlapped neither a CNS nor a mark. Statistical significance was then assessed using the chi.square function in R.

To assess statistical significance of overlaps between CNSs and ATAC-seq peaks, the proportions of CNSs fully overlapping peaks, partially overlapping peaks, and not overlapping peaks were calculated and compared to 1000 random permutations of CNS positions within Conservatory-accessible space. The locations of CNSs were randomized using bedtools shuffle, providing a BED file of Conservatory-accessible background space and allowing overlaps.

### Gene ontology enrichment

Gene ontology enrichment was performed using the clusterProfiler R package (*138*). For enrichments in each species, the corresponding annotation was retrieved from AnnotationHub (*139*). Enrichments were carried out using genes with at least one associated CNS as the background genes, compared to genes that were associated with angiosperm or tracheophyte CNSs.

### Plant materials and growth conditions

Seeds *Solanum lycopersicum* cv. M82 (LA3475), and the *SlWOX9Pro-Reg1-TFBS* (Solyc02g077390) mutants, in M82 background were our own stocks. Unless otherwise indicated, seeds were directly sown into soil in 96-cell plastic flats and grown in a greenhouse under long-day conditions (16-hr light/8-hr dark) supplemented with artificial light from high-pressure sodium bulbs (∼250 µmol m−2 s−1). The temperature ranged between 26-28°C during the day to 18-20°C during the night, with a relative humidity of 40-60%. One-month old seedlings were either transplanted to the fields at Cold Spring Harbor Laboratory (CSHL), Cold Spring Harbor, NY, or transplanted to a 4L pot and grown in the greenhouse at the same conditions described above. Plants were grown under drip irrigation and standard fertilizer regimens.

### Plant phenotyping

We defined vegetative trait phenotypes as shoot apical meristem (SAM) termination, cotyledon defects (multiple cotyledons, lobed cotyledons, and fused cotyledons), leaf fusion, and phyllotaxis defects. Vegetative trait phenotyping was done on young seedlings at various stages of vegetative growth. In order to determine if a mutant allele was embryonic lethal, an F2 population of at least 125 seedlings was screened and genotyped.

### CRISPR-Cas9 mutagenesis, plant transformation, and selection of mutant alleles

CRISPR-Cas9 mutagenesis and generation of transgenic tomato plants were performed as previously described (*140*). Briefly, guide RNAs (gRNAs) were designed using the Geneious Prime software (https://www.geneious.com) (**table S13**). The Golden Gate cloning system was used to assemble the binary vector containing the Cas9 and the specific gRNAs (*141–143*). Final binary vectors were then transformed into the tomato cultivar M82 or groundcherry by Agrobacterium tumefaciens-mediated transformation through tissue culture (*140*). First-generation transgenic plants (T_0_) were genotyped with specific primers surrounding the target sites. To generate new alleles and purify them from potential spontaneous mutations or CRISPR-Cas9 off-target effects following plant transformation, all CRISPR-Cas9 T_0_ transgenic lines were backcrossed to parental wild type M82 cultivar plants. These F1 populations were then screened for plants lacking the Cas9 transgene, and PCR products of the targeted regions were sequenced to confirm inheritance of alleles. Selected F1 plants were self-fertilized to generate F2 populations, and these segregating populations were used to validate the phenotypic effects of each allele by co-segregation. F2 or F3 homozygous mutant plants were then used for quantitative phenotypic analyses. Screening for the embryonic lethal phenotype and validation of expected segregation ratios was performed on F2 families.

### Gene tree inference

To reconstruct the evolutionary history of CNSs associated with *TB1* or *WUS*, orthologs, the predicted peptide sequences of each genome were clustered into orthogroups with OrthoFinder (*108*). Genes in the same orthogroup as the gene of interest (either *TB1* or *WUS*) that shared at least one CNS with the focal gene were retained for further analysis. For each filtered orthogroup, amino acid translations were aligned using MAFFT (version 7.526) (*76*) and trimmed with clipkit (*77*). For the *WUS* tree, only homologs from the Asteraceae and outgroup taxa were retained. Maximum likelihood trees were inferred with IQtree2 (*110*) with 1000 ultrafast bootstrap replicates (*144*). CNS presence-absence tip states were visualized with the R packages ggtree (*145*) and treeio (*146*), and CNS microsynteny was visualized with the R package gggenomes (*147*). CNSs were extracted using genome-specific GFF3 annotations with samtools (version 1.19) (*148*), aligned with MAFFT (v7.508) and visualized in Jalview (v2.11.5.0) (*149*).

### Chromatin interactions

Maize Hi-C and Hi-ChIP loop edges in B73 version 4 coordinates were obtained from previous studies (*56*, *58*) and converted to B73v5 coordinates using CrossMap v0.7.3 (*119*). *Solanum lycopersicum* cv. M82 *anantha (an)* mutants were grown in pots under greenhouse conditions (*150*). The cauliflower-like tissue from young *an* inflorescences was harvested 10 weeks after sowing, with tissue collected from two different plants to obtain two biological replicates (3.3 g and 3.1 g, respectively). Hi-C data was generated using the Arima Hi-C 2.0 Kit (PN A410110) according to the manufacturer’s user guide for plant tissues (PN A160163 v00). Nuclei extraction was done using the CelLytic Plant Nuclei Isolation/Extraction Kit according to the manufacturer’s recommendations for cell lysis and semi-pure preparation of nuclei, with the addition of a Dounce homogenizer to increase nuclei yield (Sigma, CELLYTPN1). DNA fragmentation was performed with the Bioruptor Pico (Diagenode) using 4 cycles of 15 s sonication followed by 90 s rest. Library preparation was performed using the Swift Biosciences Accel-NGS 2S Plus DNA Library Kit (PN A410110), Swift Biosciences Indexing Kit (Catalog No. 26148), KAPA Library Quantification Kit for Illumina Platforms (Catalog No. KK4824), and KAPA Library Amplification Kit (Catalog No. KK2621) according to the Arima Hi-C 2.0 library preparation guide (PN A160164 v00). Each library was sequenced in paired-end mode (2 × 250 bp) on an Illumina NovaSeq 6000 SP flowcell, yielding 463,915,437 reads for replicate 1 and 461,940,842 reads for replicate 2. Raw sequencing reads were processed using HiC-Pro v3.1.0, mapping reads to the M82 genome (SollycM82_v1.0) with a MAPQ threshold >30 (*151*, *152*). Valid read pairs were identified after filtering out invalid pairs and PCR duplicates, and both raw and normalized contact maps were generated at multiple bin sizes (5 kb, 10 kb, 20 kb, 40 kb, 150 kb, 500 kb, and 1 Mb). Following filtering, 167,352,522 and 175,494,920 valid reads were retained for replicates 1 and 2, respectively. Contact matrices were converted to .mcool format with Cooler, and the reproducibility between replicates was evaluated using HiCRep (*153*, *154*). HiC-Pro v3.1.0 was then used to merge valid read pairs from both replicates and generate contact matrices. FitHiC2 was used to identify significant chromatin looping interactions at 5 kb resolution on the merged data (q-value 0.01, maximum loop distance of 1 Mb, minimum loop distance of 10 kb), identifying a total of 9,573 loops (*155*). Loop edge sequences were then aligned to the SL4.0 genome using minimap2 with asm10 parameters (*156*) and filtered to retain cases where both edges mapped to the same SL4.0 chromosome exactly once, with primary alignment tags (tp:A:P) for both edges.

To test for CNS enrichment in loops that connected intergenic regions to genes and proximal promoters, we intersected loop edges with genes including 500 bp of upstream sequence and with CNSs. We then identified loops for which one or both edges did not intersect a gene and its 500 bp proximal promoter. For loops that connected intergenic regions, one edge was chosen at random. Loops were filtered such that the intergenic edges considered were no more than 100 kb upstream or 20 kb downstream of genes in tomato and maize, or no more than 11 kb upstream and 3 kb downstream of genes in Arabidopsis. We then constructed a 2x2 contingency table, stratifying loops by their connectivity (intergenic-genic or intergenic-intergenic) and whether the intergenic edge contained a CNS, and tested for statistical significance with a chi-square test. To evaluate the concordance of CNS-gene associations with chromatin loops in Arabidopsis, tomato and maize, we filtered to retain intergenic-genic loops that overlapped with CNSs at intergenic edges and that spanned less than the maximum upstream distance searched by Conservatory in each genome (100 kb in maize and tomato and 11 kb in Arabidopsis). For each of these loops, we then examined the gene associations of each CNS within the intergenic loop edge. If a CNS within a loop was associated with a gene, and the genic edge of that loop overlapped with the same gene, or with 500 bp of upstream sequence, it was counted as supported. All intersection operations were carried out in R using the valr package (*157*).

### Comparison of CNS space between paralog pairs

Paralog pairs were identified in each genome using DupGenFinder (*158*) with *Amborella trichopoda* as the outgroup. For each paralog pair, the normalized number of CNSs specific to one pair was compared to the total CNS associated with both pairs.

## Acknowledgments

We thank the members of the Bartlett, Lippman, and Efroni laboratories for their support and feedback; T. Harrington, G. Robitaille, B. Seman for technical assistance; T. Mulligan, K. Schlecht, S. Qiao, and B. Fitzgerald for assistance with plant care; H. Golan for website design.

## Funding

Binational Science Foundation 2021660 (IE)

Israeli Science Foundation grant 928/22 (IE)

Howard Hughes Medical Institute International Research Scholar grant 55008730 (IE)

National Science Foundation Plant Genome Research Program grant IOS-2129189 (DJ, MEB, ZBL)

USDA-NIFA Postdoctoral Fellowship (KRA) Gatsby Foundation (MEB)

Howard Hughes Medical Institute (ZBL)

## Author contributions

Conceptualization: DJ, MEB, ZBL, IE

Investigation: KRA, AH, DC, HY, AEdN, ST, AS, MEB, ZBL, IE

Software: AH, AS, IE

Funding acquisition: DJ, MEB, ZBL, IE Supervision: MEB, ZBL, IE

Writing – original draft: KRA, AH, MEB, ZBL, IE

Writing – reviewing and editing: KRA, AH, DJ, MEB, ZBL, IE

## Competing interests

Z.B.L and D.J. are consultants for Inari Agriculture, and Z.B.L is a member of their scientific strategy board. All other authors declare that they have no competing interests.

## Data and materials availability

Raw and processed data for Conservatory are available from www.conservatorycns.com and Zenodo (*29*); Conservatory code is available at https://github.com/idanefroni/Conservatory and <u>doi.org/10.5281/zenodo.18478543</u>. Tomato Hi-C and ATAC-Seq raw data are available from NCBI Gene Expression Omnibus record GSE307325. All reasonable requests for biological material will be fulfilled by the corresponding authors.

## Supplementary Materials

**Fig. S1.**
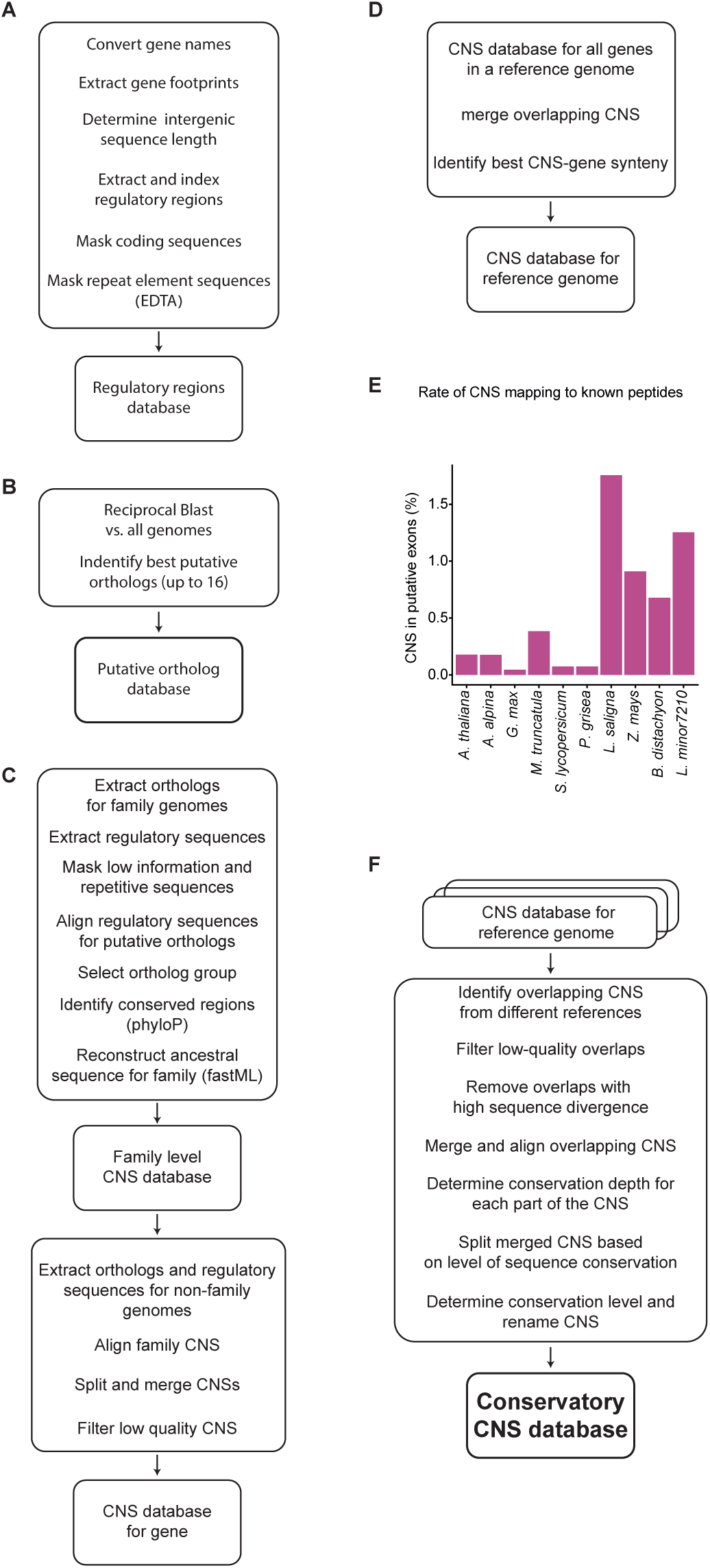
Flow chart of the Conservatory algorithm. (**A**) Preprocessing of genome sequence and annotation data. (**B**) Construction of the homologs database. (**C**) Identification of CNS per gene. (**D**) Merging of CNS for all genes in a given reference genome. (**E**) Mapping rate of Identified CNSs to known peptides for the 10 referenced genomes. (**F**) Merging CNS databases from different reference genomes.

**Fig. S2.**
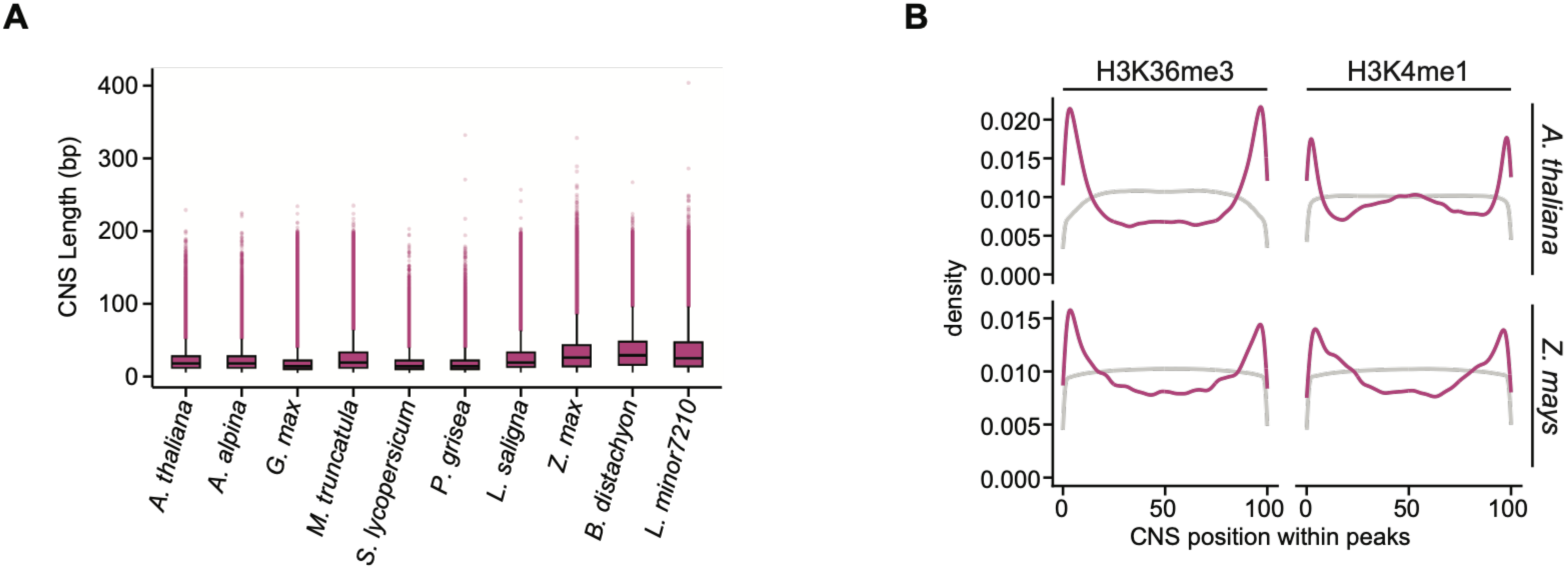
CNS lengths and positions with regions of activating chromatin marks. **(A)** Distributions of CNS lengths in 10 reference genomes. **(B)** Distribution of CNS (pink) and gene coding regions (gray) positions within regions of activating chromatin marks. CNS are preferentially located at the edge of these regions, suggesting they overlap due to their proximity to genes.

**Fig. S3.**
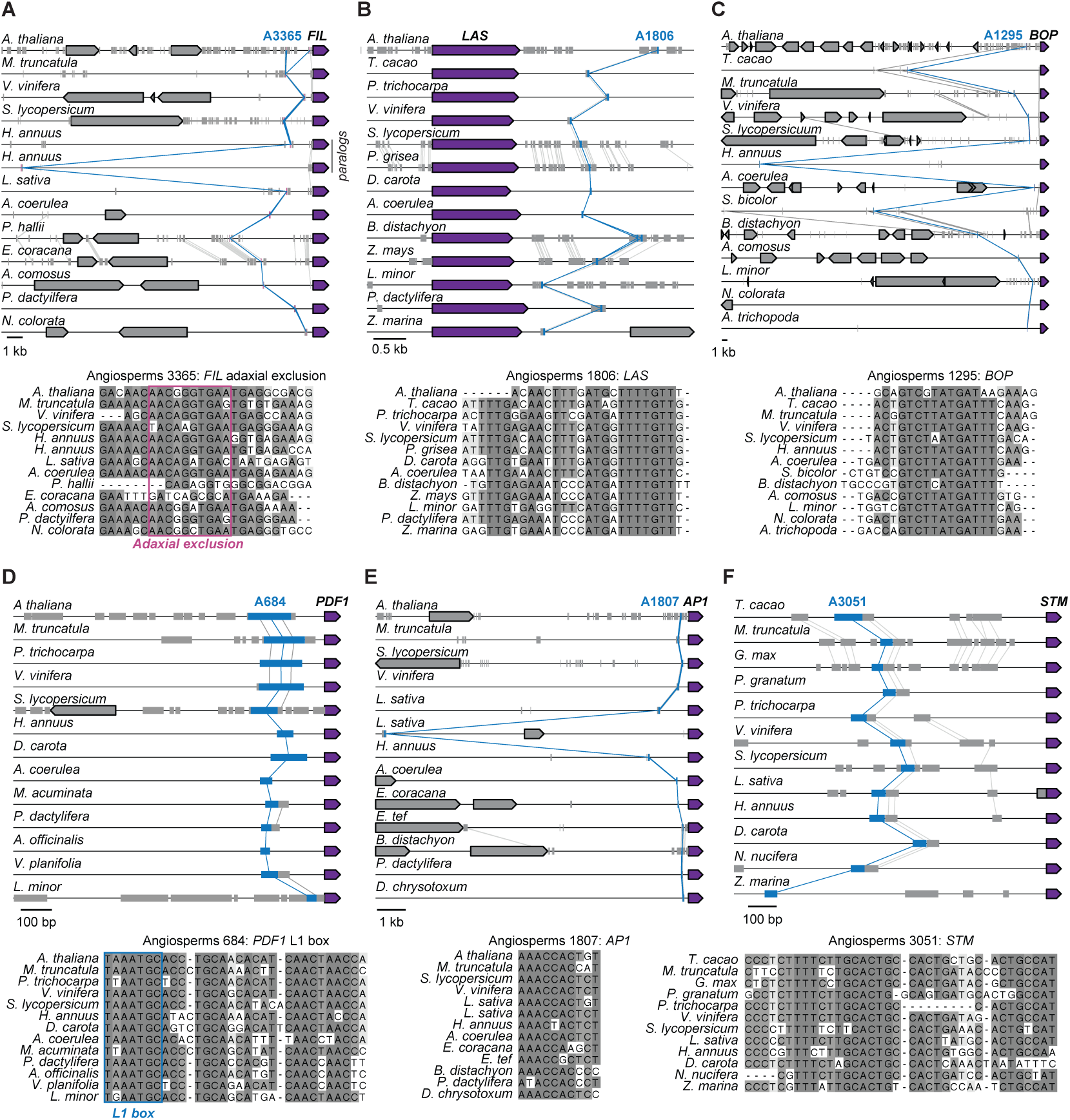
CNSs identified within experimentally validated CREs. (A-F) Microsynteny maps and multiple sequence alignments for CNSs located within experimentally validated CREs of *FILAMENTOUS FLOWER (FIL)* (*37*) (A), *LATERAL SUPPRESSOR (LAS)* (*36*) (B), *BLADE-ON-PETIOLE (BOP)* (*13*) (C), *PROTODERMAL FACTOR 1 (PDF1)* (*39*) (D), *APETALA1 (AP1)* (*38*) (E), and *SHOOT MERISTEMLESS (STM)* (*35*, *159*) (F). See **table S7** for gene names.

**Fig. S4.**
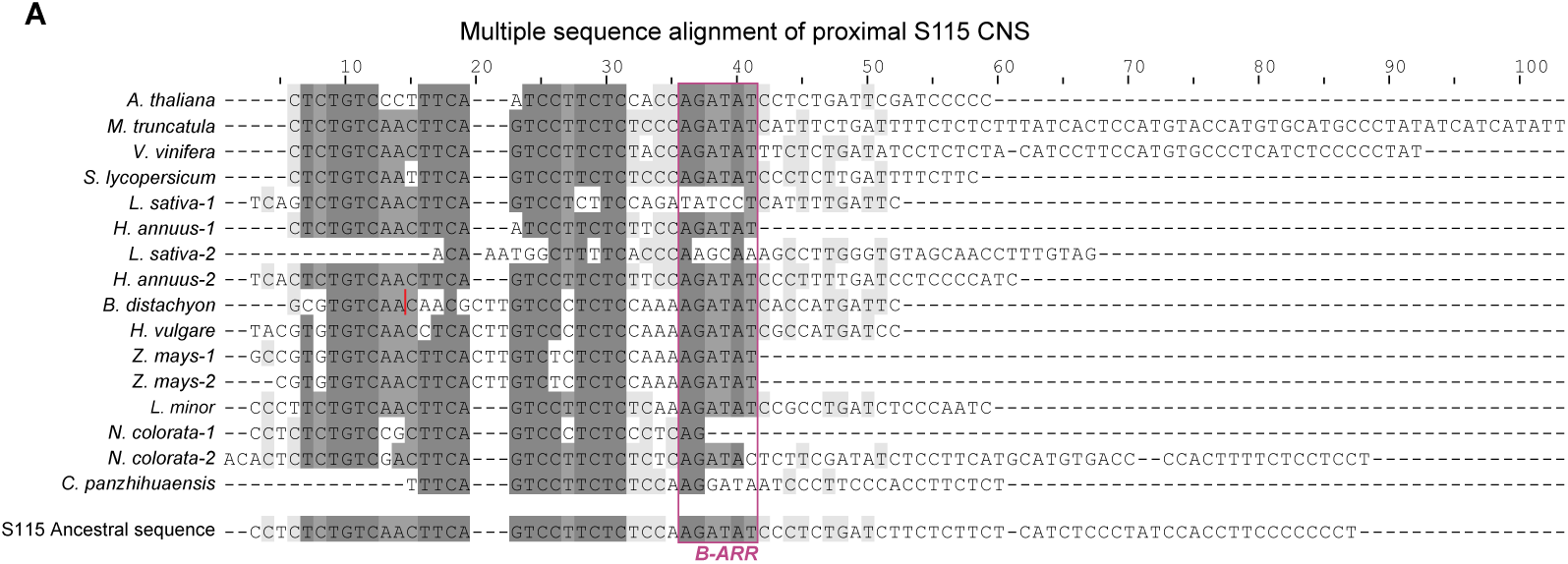
Multiple alignment of the S115 seed plant CNS. The B-ARR binding site is marked by the pink box. See **table S7** for gene names.

**Fig. S5.**
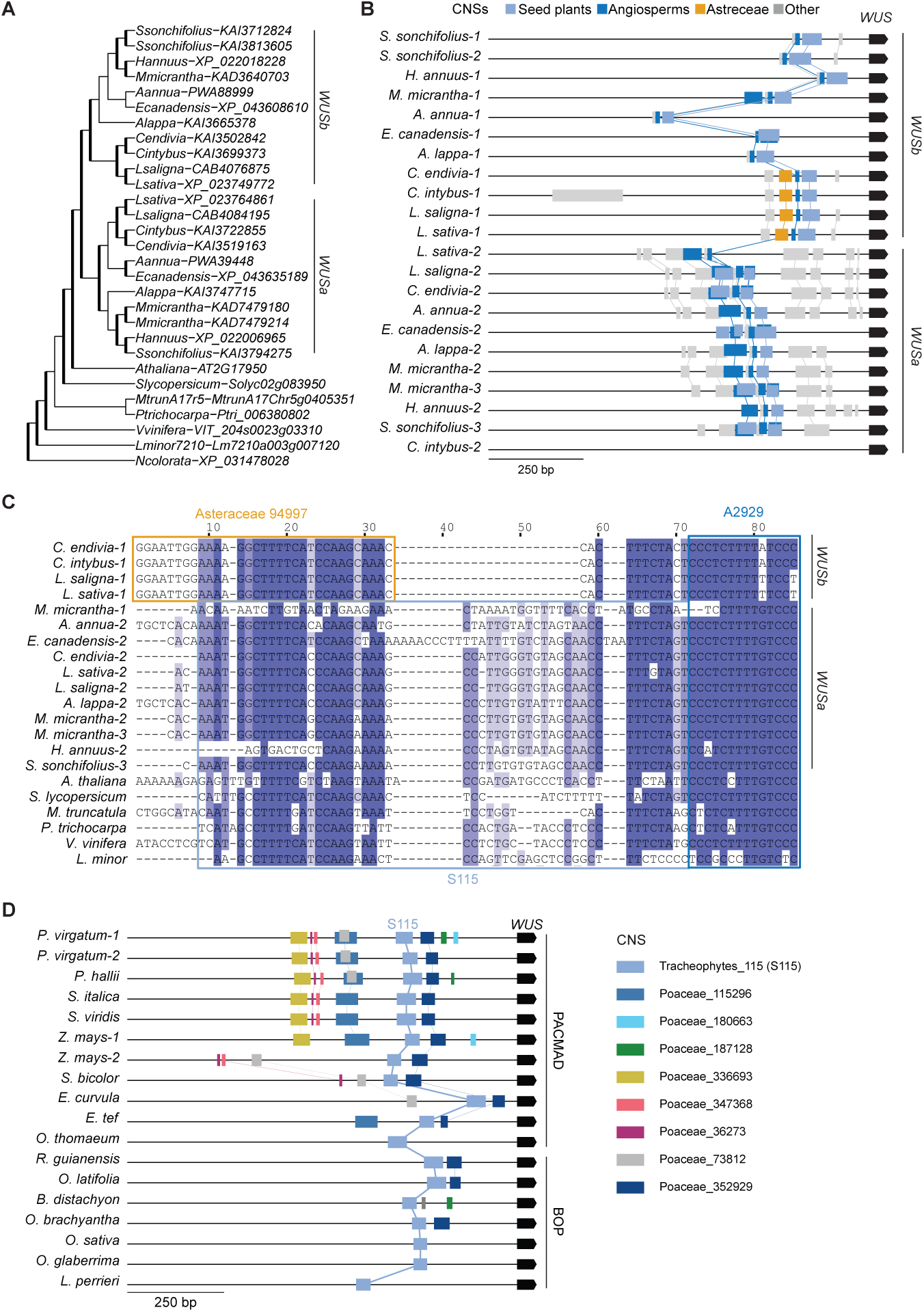
Gene duplication results in paralog-specific divergence of ancient CNSs. **(A)** Gene tree of WUS showing the duplication at the base of the Asteraceae that gave rise to *WUSa* and *WUS*b. **(B)** Microsynteny map of *WUSa* and *WUSb* in the Asteraceae. Note that *WUSa* maintains greater conservation of the meristem CNS module, while *WUSb* exhibits increased divergence with Asteraceae-specific CNSs (shown in orange). **(C)** Multiple sequence alignment of the ancient S115 from representative angiosperms and the collinear Asteraceae-specific CNS in *WUSa*. **(D)** Microsynteny map of *WUS* homologs in the grasses, showing conservation among paralogs within the PACMAD clade. See **table S7** for gene names.

**Fig. S6.**
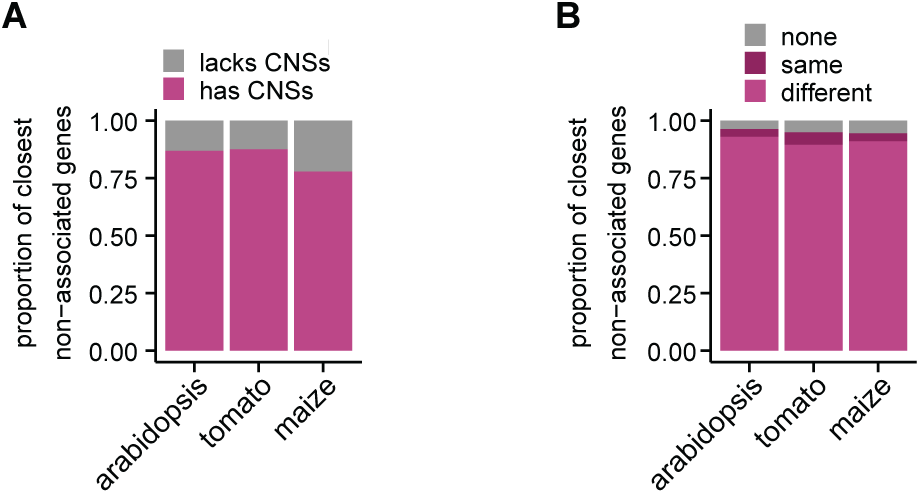
Association of CNS with non-adjacent genes. (**A**) Proportion of genes located between a CNS and an associated gene that have CNS of their own. (**B**) Proportion of genes located between CNS and an associated gene that belong to either the same or different orthogroup. The vast majority belong to a different orthogroup, indicating that the gene skipping is not due to tandem duplications.

## Notes

https://www.conservatorycns.com

### Summary of Updates

Updates and clarifications to the text. Additional details added for supplemental tables.

## References and Notes

1. S. B. Carroll, Evo-devo and an expanding evolutionary synthesis: a genetic theory of morphological evolution. Cell 134, 25–36 (2008).

2. P. J. Wittkopp, G. Kalay, Cis-regulatory elements: molecular mechanisms and evolutionary processes underlying divergence. Nat. Rev. Genet. 13, 59–69 (2011).

3. D. L. Stern, V. Orgogozo, The loci of evolution: how predictable is genetic evolution? Evolution 62, 2155–2177 (2008).

4. D. Villar, P. Flicek, D. T. Odom, Evolution of transcription factor binding in metazoans - mechanisms and functional implications. Nat. Rev. Genet. 15, 221–233 (2014).

5. X. Li, R. J. Schmitz, Cis-regulatory dynamics in plant domestication. Trends Genet., doi: 10.1016/j.tig.2025.02.005 (2025).

6. A. P. Marand, L. Jiang, F. Gomez-Cano, M. A. A. Minow, X. Zhang, J. P. Mendieta, Z. Luo, S. Bang, H. Yan, C. Meyer, L. Schlegel, F. Johannes, R. J. Schmitz, The genetic architecture of cell type-specific cis regulation in maize. Science 388, eads6601 (2025).

7. D. Villar, C. Berthelot, S. Aldridge, T. F. Rayner, M. Lukk, M. Pignatelli, T. J. Park, R. Deaville, J. T. Erichsen, A. J. Jasinska, J. M. A. Turner, M. F. Bertelsen, E. P. Murchison, P. Flicek, D. T. Odom, Enhancer evolution across 20 mammalian species. Cell 160, 554–566 (2015).

8. E. Z. Kvon, O. K. Kamneva, U. S. Melo, I. Barozzi, M. Osterwalder, B. J. Mannion, V. Tissières, C. S. Pickle, I. Plajzer-Frick, E. A. Lee, M. Kato, T. H. Garvin, J. A. Akiyama, V. Afzal, J. Lopez-Rios, E. M. Rubin, D. E. Dickel, L. A. Pennacchio, A. Visel, Progressive loss of function in a limb enhancer during snake evolution. Cell 167, 633–642.e11 (2016).

9. A. Hendelman, S. Zebell, D. Rodriguez-Leal, N. Dukler, G. Robitaille, X. Wu, J. Kostyun, L. Tal, P. Wang, M. E. Bartlett, Y. Eshed, I. Efroni, Z. B. Lippman, Conserved pleiotropy of an ancient plant homeobox gene uncovered by cis-regulatory dissection. Cell 184, 1724–1739.e16 (2021).

10. A. Lanctot, A. Hendelman, P. Udilovich, G. M. Robitaille, Z. B. Lippman, Antagonizing cis-regulatory elements of a conserved flowering gene mediate developmental robustness. Proc. Natl. Acad. Sci. U. S. A. 122, e2421990122 (2025).

11. D. Ciren, S. Zebell, Z. B. Lippman, Extreme restructuring of cis-regulatory regions controlling a deeply conserved plant stem cell regulator. PLoS Genet. 20, e1011174 (2024).

12. X. Wang, L. Aguirre, D. Rodríguez-Leal, A. Hendelman, M. Benoit, Z. B. Lippman, Dissecting cis-regulatory control of quantitative trait variation in a plant stem cell circuit. Nat. Plants 7, 419–427 (2021).

13. T. C. Tran, K. Mähl, C. Kappel, Y. Dakhiya, A. Sampathkumar, A. Sicard, M. Lenhard, Altered interactions between cis-regulatory elements partially resolve BLADE-ON-PETIOLE genetic redundancy in Capsella rubella. Plant Cell 36, 4637–4657 (2024).

14. S. G. Zebell, C. Martí-Gómez, B. Fitzgerald, C. P. Cunha, M. Lach, B. M. Seman, Hendelman, S. Sretenovic, Y. Qi, M. Bartlett, Y. Eshed, D. M. McCandlish, Z. B. Lippman, Cryptic variation fuels plant phenotypic change through hierarchical epistasis. Nature 644, 984–992 (2025).

15. M. J. Christmas, I. M. Kaplow, D. P. Genereux, M. X. Dong, G. M. Hughes, X. Li, P. F. Sullivan, A. G. Hindle, G. Andrews, J. C. Armstrong, M. Bianchi, A. M. Breit, M. Diekhans, C. Fanter, N. M. Foley, D. B. Goodman, L. Goodman, K. C. Keough, B. Kirilenko, A. Kowalczyk, C. Lawless, A. L. Lind, J. R. S. Meadows, L. R. Moreira, R. W. Redlich, L. Ryan, R. Swofford, A. Valenzuela, F. Wagner, O. Wallerman, A. R. Brown, J. Damas, K. Fan, J. Gatesy, J. Grimshaw, J. Johnson, S. V. Kozyrev, A. J. Lawler, V. D. Marinescu, K. M. Morrill, A. Osmanski, N. S. Paulat, B. N. Phan, S. K. Reilly, D. E. Schäffer, C. Steiner, M. A. Supple, A. P. Wilder, M. E. Wirthlin, J. R. Xue, Zoonomia Consortium§, B. W. Birren, S. Gazal, R. M. Hubley, K.-P. Koepfli, T. Marques-Bonet, W. K. Meyer, M. Nweeia, P. C. Sabeti, B. Shapiro, A. F. A. Smit, M. S. Springer, E. C. Teeling, Z. Weng, M. Hiller, D. L. Levesque, H. A. Lewin, W. J. Murphy, A. Navarro, B. Paten, K. S. Pollard, D. A. Ray, I. Ruf, O. A. Ryder, A. R. Pfenning, K. Lindblad-Toh, E. K. Karlsson, Evolutionary constraint and innovation across hundreds of placental mammals. Science 380, eabn3943 (2023).

16. J. Stiller, S. Feng, A.-A. Chowdhury, I. Rivas-González, D. A. Duchêne, Q. Fang, Y. Deng, A. Kozlov, A. Stamatakis, S. Claramunt, J. M. T. Nguyen, S. Y. W. Ho, B. C. Faircloth, J. Haag, P. Houde, J. Cracraft, M. Balaban, U. Mai, G. Chen, R. Gao, C. Zhou, Y. Xie, Z. Huang, Z. Cao, Z. Yan, H. A. Ogilvie, L. Nakhleh, B. Lindow, B. Morel, J. Fjeldså, P. A. Hosner, R. R. da Fonseca, B. Petersen, J. A. Tobias, T. Székely, J. D. Kennedy, A. H. Reeve, A. Liker, M. Stervander, A. Antunes, D. T. Tietze, M. F. Bertelsen, F. Lei, C. Rahbek, G. R. Graves, M. H. Schierup, T. Warnow, E. L. Braun, M. T. P. Gilbert, E. D. Jarvis, S. Mirarab, G. Zhang, Complexity of avian evolution revealed by family-level genomes. Nature 629, 851–860 (2024).

17. E. S. Wong, D. Zheng, S. Z. Tan, N. L. Bower, V. Garside, G. Vanwalleghem, F. Gaiti, E. Scott, B. M. Hogan, K. Kikuchi, E. McGlinn, M. Francois, B. M. Degnan, Deep conservation of the enhancer regulatory code in animals. Science 370, eaax8137 (2020).

18. M. H. Q. Phan, T. Zehnder, F. Puntieri, A. Magg, B. Majchrzycka, M. Antonović, H. Wieler, B.-W. Lo, D. Baranasic, B. Lenhard, F. Müller, M. Vingron, D. M. Ibrahim, Conservation of regulatory elements with highly diverged sequences across large evolutionary distances. Nat. Genet. 57, 1524–1534 (2025).

19. I. Braasch, A. R. Gehrke, J. J. Smith, K. Kawasaki, T. Manousaki, J. Pasquier, A. Amores, T. Desvignes, P. Batzel, J. Catchen, A. M. Berlin, M. S. Campbell, D. Barrell, K. J. Martin, J. F. Mulley, V. Ravi, A. P. Lee, T. Nakamura, D. Chalopin, S. Fan, D. Wcisel, C. Cañestro, J. Sydes, F. E. G. Beaudry, Y. Sun, J. Hertel, M. J. Beam, M. Fasold, M. Ishiyama, J. Johnson, S. Kehr, M. Lara, J. H. Letaw, G. W. Litman, R. T. Litman, M. Mikami, T. Ota, N. R. Saha, L. Williams, P. F. Stadler, H. Wang, J. S. Taylor, Q. Fontenot, A. Ferrara, S. M. J. Searle, B. Aken, M. Yandell, I. Schneider, J. A. Yoder, J.-N. Volff, A. Meyer, C. T. Amemiya, B. Venkatesh, P. W. H. Holland, Y. Guiguen, J. Bobe, N. H. Shubin, F. Di Palma, J. Alföldi, K. Lindblad-Toh, J. H. Postlethwait, The spotted gar genome illuminates vertebrate evolution and facilitates human-teleost comparisons. Nat. Genet. 48, 427–437 (2016).

20. B. Song, E. S. Buckler, M. C. Stitzer, New whole-genome alignment tools are needed for tapping into plant diversity. Trends Plant Sci. 29, 355–369 (2024).

21. A. Haudry, A. E. Platts, E. Vello, D. R. Hoen, M. Leclercq, R. J. Williamson, E. Forczek, Z. Joly-Lopez, J. G. Steffen, K. M. Hazzouri, K. Dewar, J. R. Stinchcombe, D. J. Schoen, X. Wang, J. Schmutz, C. D. Town, P. P. Edger, J. C. Pires, K. S. Schumaker, D. E. Jarvis, T. Mandáková, M. A. Lysak, E. van den Bergh, M. E. Schranz, P. M. Harrison, A. M. Moses, T. E. Bureau, S. I. Wright, M. Blanchette, An atlas of over 90,000 conserved noncoding sequences provides insight into crucifer regulatory regions. Nat. Genet. 45, 891–898 (2013).

22. B. Song, E. S. Buckler, H. Wang, Y. Wu, E. Rees, E. A. Kellogg, D. J. Gates, M. Khaipho-Burch, P. J. Bradbury, J. Ross-Ibarra, M. B. Hufford, M. C. Romay, Conserved noncoding sequences provide insights into regulatory sequence and loss of gene expression in maize. Genome Res. 31, 1245–1257 (2021).

23. M. C. Stitzer, A. S. Seetharam, A. Scheben, S.-K. Hsu, A. J. Schulz, T. AuBuchon-Elder, M. El-Walid, T. H. Ferebee, C. O. Hale, T. La, Z.-Y. Liu, S. J. McMorrow, P. Minx, A. Phillips, M. Syring, T. Wrightsman, J. Zhai, R. Pasquet, C. McAllister, S. Malcomber, P. Traiperm, D. Layton, J. Zhong, D. E. Costich, R. K. Dawe, K. Fengler, C. Harris, Z. Irelan, V. Llaca, P. Parakkal, G. Zastrow-Hayes, M. R. Woodhouse, E. K. S. Cannon, J. Portwood II, C. M. Andorf, P. S. Albert, J. A. Birchler, A. Siepel, J. Ross-Ibarra, M. C. Romay, E. Kellogg, E. S. Buckler IV, M. Hufford, Extensive genome evolution distinguishes maize within a stable tribe of grasses, bioRxiv (2025). 10.1101/2025.01.22.633974.

24. H. Xin, X. Liu, S. Chai, X. Yang, H. Li, B. Wang, Y. Xu, S. Lin, X. Zhong, B. Liu, Z. Lu, Z. Zhang, Identification and functional characterization of conserved cis-regulatory elements responsible for early fruit development in cucurbit crops. Plant Cell 36, 2272–2288 (2024).

25. X. Lai, S. Behera, Z. Liang, Y. Lu, J. S. Deogun, J. C. Schnable, STAG-CNS: An order-aware conserved noncoding sequences discovery tool for arbitrary numbers of species. Mol. Plant 10, 990–999 (2017).

26. S. Magallón, S. Gómez-Acevedo, L. L. Sánchez-Reyes, T. Hernández-Hernández, A metacalibrated time-tree documents the early rise of flowering plant phylogenetic diversity. New Phytol. 207, 437–453 (2015).

27. K. S. Pollard, M. J. Hubisz, K. R. Rosenbloom, A. Siepel, Detection of nonneutral substitution rates on mammalian phylogenies. Genome Res. 20, 110–121 (2010).

28. H. Ashkenazy, O. Penn, A. Doron-Faigenboim, O. Cohen, G. Cannarozzi, O. Zomer, T. Pupko, FastML: a web server for probabilistic reconstruction of ancestral sequences. Nucleic Acids Res. 40, W580–4 (2012).

29. I. Efroni, Conservatory - Conserved non-coding sequences in plants, Zenodo (2026); 10.5281/ZENODO.18484694.

30. J. Van de Velde, M. Van Bel, D. Vaneechoutte, K. Vandepoele, A collection of conserved noncoding sequences to study gene regulation in flowering plants. Plant Physiol. 171, 2586–2598 (2016).

31. D. Burgess, M. Freeling, The most deeply conserved noncoding sequences in plants serve similar functions to those in vertebrates despite large differences in evolutionary rates. Plant Cell 26, 946–961 (2014).

32. Z. Lu, A. P. Marand, W. A. Ricci, C. L. Ethridge, X. Zhang, R. J. Schmitz, The prevalence, evolution and chromatin signatures of plant regulatory elements. Nat. Plants 5, 1250–1259 (2019).

33. J. Candela-Ferre, B. Diego-Martin, J. Pérez-Alemany, J. Gallego-Bartolomé, Mind the gap: Epigenetic regulation of chromatin accessibility in plants. Plant Physiol. 194, 1998–2016 (2024).

34. S. Ramírez-Barahona, H. Sauquet, S. Magallón, The delayed and geographically heterogeneous diversification of flowering plant families. *Nat*. Ecol. Evol. 4, 1232–1238 (2020).

35. H. Breuninger, E. Rikirsch, M. Hermann, M. Ueda, T. Laux, Differential expression of WOX genes mediates apical-basal axis formation in the Arabidopsis embryo. Dev. Cell 14, 867–876 (2008).

36. J. A. Aguilar-Martínez, N. Uchida, B. Townsley, D. A. West, A. Yanez, N. Lynn, S. Kimura, N. Sinha, Transcriptional, posttranscriptional, and posttranslational regulation of SHOOT MERISTEMLESS gene expression in Arabidopsis determines gene function in the shoot apex. Plant Physiol. 167, 424–442 (2015).

37. B. Raatz, A. Eicker, G. Schmitz, E. Fuss, D. Müller, S. Rossmann, K. Theres, Specific expression of LATERAL SUPPRESSOR is controlled by an evolutionarily conserved 3’ enhancer: LAS/Ls is regulated by a downstream enhancer. Plant J. 68, 400–412 (2011).

38. K. Watanabe, K. Okada, Two discrete cis elements control the Abaxial side-specific expression of the FILAMENTOUS FLOWER gene in Arabidopsis. Plant Cell 15, 2592–2602 (2003).

39. L. Ye, B. Wang, W. Zhang, H. Shan, H. Kong, Gains and losses of Cis-regulatory elements led to divergence of the Arabidopsis APETALA1 and CAULIFLOWER duplicate genes in the time, space, and level of expression and regulation of one paralog by the other. Plant Physiol. 171, 1055–1069 (2016).

40. M. Abe, T. Takahashi, Y. Komeda, Identification of a cis-regulatory element for L1 layer-specific gene expression, which is targeted by an L1-specific homeodomain protein: Identification of the L1 box. Plant J. 26, 487–494 (2001).

41. M. Kerstens, Y. Boele, A. Moralez-Cruz, C. Roelofsen, P. Wang, L. A. Baumgart, R. O’Malley, G. Sanchez-Perez, B. Scheres, V. Willemsen, Two deeply conserved non-coding sequences control PLETHORA1/2 expression and coordinate embryo and root development. Plant Commun. 0, 101466 (2025).

42. M. Somssich, B. I. Je, R. Simon, D. Jackson, CLAVATA-WUSCHEL signaling in the shoot meristem. Development 143, 3238–3248 (2016).

43. I. Bäurle, T. Laux, Regulation of WUSCHEL transcription in the stem cell niche of the Arabidopsis shoot meristem. Plant Cell 17, 2271–2280 (2005).

44. T.-Q. Zhang, H. Lian, C.-M. Zhou, L. Xu, Y. Jiao, J.-W. Wang, A two-step model for de Novo activation of WUSCHEL during plant shoot regeneration. Plant Cell 29, 1073–1087 (2017).

45. Z. Chen, W. Li, C. Gaines, A. Buck, M. Galli, A. Gallavotti, Structural variation at the maize WUSCHEL1 locus alters stem cell organization in inflorescences. Nat. Commun. 12, 2378 (2021).

46. J. Nardmann, W. Werr, The shoot stem cell niche in angiosperms: expression patterns of WUS orthologues in rice and maize imply major modifications in the course of mono- and dicot evolution. Mol. Biol. Evol. 23, 2492–2504 (2006).

47. B. I. Je, J. Gruel, Y. K. Lee, P. Bommert, E. D. Arevalo, A. L. Eveland, Q. Wu, Goldshmidt, R. Meeley, M. Bartlett, M. Komatsu, H. Sakai, H. Jönsson, D. Jackson, Signaling from maize organ primordia via FASCIATED EAR3 regulates stem cell proliferation and yield traits. Nat. Genet. 48, 785–791 (2016).

48. T. H. Kebrom, B. L. Burson, S. A. Finlayson, Phytochrome B represses Teosinte Branched1 expression and induces sorghum axillary bud outgrowth in response to light signals. Plant Physiol. 140, 1109–1117 (2006).

49. J. A. Aguilar-Martínez, C. Poza-Carrión, P. Cubas, Arabidopsis BRANCHED1 acts as an integrator of branching signals within axillary buds. Plant Cell 19, 458–472 (2007).

50. J. Doebley, A. Stec, L. Hubbard, The evolution of apical dominance in maize. Nature 386, 485–488 (1997).

51. R. M. Clark, E. Linton, J. Messing, J. F. Doebley, Pattern of diversity in the genomic region near the maize domestication gene tb1. Proc. Natl. Acad. Sci. U. S. A. 101, 700–707 (2004).

52. M.-S. Remigereau, G. Lakis, S. Rekima, M. Leveugle, M. C. Fontaine, T. Langin, Sarr, T. Robert, Cereal domestication and evolution of branching: evidence for soft selection in the Tb1 orthologue of pearl millet (Pennisetum glaucum [L.] R. Br.). PLoS One 6, e22404 (2011).

53. X. Wu, Y. Liu, H. Luo, L. Shang, C. Leng, Z. Liu, Z. Li, X. Lu, H. Cai, H. Hao, H.-C. Jing, Genomic footprints of sorghum domestication and breeding selection for multiple end uses. Mol. Plant 15, 537–551 (2022).

54. A. Studer, Q. Zhao, J. Ross-Ibarra, J. Doebley, Identification of a functional transposon insertion in the maize domestication gene tb1. Nat. Genet. 43, 1160–1163 (2011).

55. A. Sanyal, B. R. Lajoie, G. Jain, J. Dekker, The long-range interaction landscape of gene promoters. Nature 489, 109–113 (2012).

56. W. A. Ricci, Z. Lu, L. Ji, A. P. Marand, C. L. Ethridge, N. G. Murphy, J. M. Noshay, M. Galli, M. K. Mejía-Guerra, M. Colomé-Tatché, F. Johannes, M. J. Rowley, V. G. Corces, J. Zhai, M. J. Scanlon, E. S. Buckler, A. Gallavotti, N. M. Springer, R. J. Schmitz, X. Zhang, Widespread long-range cis-regulatory elements in the maize genome. Nat. Plants 5, 1237–1249 (2019).

57. C. Liu, C. Wang, G. Wang, C. Becker, M. Zaidem, D. Weigel, Genome-wide analysis of chromatin packing in Arabidopsis thaliana at single-gene resolution. Genome Res. 26, 1057–1068 (2016).

58. Y. Sun, L. Dong, Y. Zhang, D. Lin, W. Xu, C. Ke, L. Han, L. Deng, G. Li, D. Jackson, X. Li, F. Yang, 3D genome architecture coordinates trans and cis regulation of differentially expressed ear and tassel genes in maize. Genome Biol. 21, 143 (2020).

59. E. A. Kellogg, Evolutionary history of the grasses. Plant Physiol. 125, 1198–1205 (2001).

60. E. A. Kellogg, Flowering Plants. Monocots: Poaceae (Springer International Publishing, Cham, Switzerland, ed. 2015, 2015)*The Families and Genera of Vascular Plants*.

61. J. M. C. McDonald, R. D. Reed, Beyond modular enhancers: new questions in cis-regulatory evolution. Trends Ecol. Evol. 39, 1035–1046 (2024).

62. D. Singh, T. Hoyt, S. V. Yi, Evolution of Enhancers through Duplication, Evolutionary Biology (2025). https://www.biorxiv.org/content/10.1101/2025.04.09.648062v1.abstract.

63. M. Benoit, K. M. Jenike, J. W. Satterlee, S. Ramakrishnan, I. Gentile, A. Hendelman, M. Passalacqua, H. Suresh, H. Shohat, G. M. Robitaille, B. Fitzgerald, M. Alonge, X. Wang, R. Santos, J. He, S. Ou, H. Golan, Y. Green, K. Swartwood, N. G. Karavolias, G. P. Sierra, A. Orejuela, F. Roda, S. Goodwin, W. R. McCombie, E. Kizito, E. Gagnon, S. Knapp, T. Särkinen, A. Frary, J. Gillis, J. Van Eck, M. C. Schatz, Z. Lippman, Solanum pan-genetics reveals paralogues as contingencies in crop engineering. Nature 640, 135–145 (2025).

64. X. Li, X. Zhang, R. J. Schmitz, Using single-cell genomics to explore transcriptional divergence and cis -regulatory dynamics of duplicated genes, Evolutionary Biology (2025). https://www.biorxiv.org/content/10.1101/2025.05.07.652760v2.full.pdf.

65. M. E. Wirthlin, T. A. Schmid, J. E. Elie, X. Zhang, A. Kowalczyk, R. Redlich, V. A. Shvareva, A. Rakuljic, M. B. Ji, N. S. Bhat, I. M. Kaplow, D. E. Schäffer, A. J. Lawler, Z. Wang, B. N. Phan, S. Annaldasula, A. R. Brown, T. Lu, B. K. Lim, E. Azim, Zoonomia Consortium, N. L. Clark, W. K. Meyer, S. L. K. Pond, M. Chikina, M. M. Yartsev, A. R. Pfenning, G. Andrews, J. C. Armstrong, M. Bianchi, B. W. Birren, K. R. Bredemeyer, A. M. Breit, M. J. Christmas, H. Clawson, J. Damas, F. Di Palma, M. Diekhans, M. X. Dong, E. Eizirik, K. Fan, C. Fanter, N. M. Foley, K. Forsberg-Nilsson, C. J. Garcia, J. Gatesy, S. Gazal, D. P. Genereux, L. Goodman, J. Grimshaw, M. K. Halsey, A. J. Harris, G. Hickey, M. Hiller, A. G. Hindle, R. M. Hubley, G. M. Hughes, J. Johnson, D. Juan, I. M. Kaplow, E. K. Karlsson, K. C. Keough, B. Kirilenko, K.-P. Koepfli, J. M. Korstian, A. Kowalczyk, S. V. Kozyrev, A. J. Lawler, C. Lawless, T. Lehmann, D. L. Levesque, H. A. Lewin, X. Li, A. Lind, K. Lindblad-Toh, A. Mackay-Smith, V. D. Marinescu, T. Marques-Bonet, V. C. Mason, J. R. S. Meadows, W. K. Meyer, J. E. Moore, L. R. Moreira, D. D. Moreno-Santillan, K. M. Morrill, G. Muntané, W. J. Murphy, A. Navarro, M. Nweeia, S. Ortmann, A. Osmanski, B. Paten, N. S. Paulat, A. R. Pfenning, B. N. Phan, K. S. Pollard, H. E. Pratt, D. A. Ray, S. K. Reilly, J. R. Rosen, I. Ruf, L. Ryan, O. A. Ryder, P. C. Sabeti, D. E. Schäffer, A. Serres, B. Shapiro, A. F. A. Smit, M. Springer, C. Srinivasan, C. Steiner, J. M. Storer, K. A. M. Sullivan, P. F. Sullivan, E. Sundström, M. A. Supple, R. Swofford, J.-E. Talbot, E. Teeling, J. Turner-Maier, A. Valenzuela, F. Wagner, O. Wallerman, C. Wang, J. Wang, Z. Weng, A. P. Wilder, M. E. Wirthlin, J. R. Xue, X. Zhang, Vocal learning-associated convergent evolution in mammalian proteins and regulatory elements. Science 383, eabn3263 (2024).

66. P. L. van der Jagt, S. Oud, R. M. A. Vroomans, Ubiquitous systems drift in the evolution of development, bioRxiv (2025). 10.1101/2025.04.23.650280.

67. Á. McColgan, J. DiFrisco, Understanding developmental system drift. Development 151, dev203054 (2024).

68. E. Ernst, B. Abramson, K. Acosta, P. T. N. Hoang, C. Mateo-Elizalde, V. Schubert, B. Pasaribu, P. S. Albert, N. Hartwick, K. Colt, A. Aylward, U. Ramu, J. A. Birchler, I. Schubert, E. Lam, T. P. Michael, R. A. Martienssen, Duckweed genomes and epigenomes underlie triploid hybridization and clonal reproduction. Curr. Biol. 35, 1828–1847.e9 (2025).

69. J. Zhai, A. Gokaslan, Y. Schiff, A. Berthel, Z.-Y. Liu, W.-Y. Lai, Z. R. Miller, A. Scheben, M. C. Stitzer, M. C. Romay, E. S. Buckler, V. Kuleshov, Cross-species modeling of plant genomes at single-nucleotide resolution using a pretrained DNA language model. Proc. Natl. Acad. Sci. U. S. A. 122, e2421738122 (2025).

70. G. Li, L. An, W. Yang, L. Yang, T. Wei, J. Shi, J. Wang, J. H. Doonan, K. Xie, R. Fernie, E. S. Lagudah, R. A. Wing, C. Gao, Integrated biotechnological and AI innovations for crop improvement. Nature 643, 925–937 (2025).

71. S. Ou, W. Su, Y. Liao, K. Chougule, J. R. A. Agda, A. J. Hellinga, C. S. B. Lugo, T. A. Elliott, D. Ware, T. Peterson, N. Jiang, C. N. Hirsch, M. B. Hufford, Benchmarking transposable element annotation methods for creation of a streamlined, comprehensive pipeline. Genome Biol. 20, 275 (2019).

72. F. Tegenfeldt, D. Kuznetsov, M. Manni, M. Berkeley, E. M. Zdobnov, E. V. Kriventseva, OrthoDB and BUSCO update: annotation of orthologs with wider sampling of genomes. Nucleic Acids Res. 53, D516–D522 (2025).

73. H. Li, Protein-to-genome alignment with miniprot. Bioinformatics 39 (2023).

74. S. R. Eddy, Accelerated profile HMM searches. PLoS Comput. Biol. 7, e1002195 (2011).

75. B. Bushnell, BBMap: A Fast, Accurate, Splice-Aware Aligner (2014). https://escholarship.org/uc/item/1h3515gn.

76. K. Katoh, D. M. Standley, MAFFT multiple sequence alignment software version 7: improvements in performance and usability. Mol. Biol. Evol. 30, 772–780 (2013).

77. J. L. Steenwyk, T. J. Buida 3rd, Y. Li, X.-X. Shen, A. Rokas, ClipKIT: A multiple sequence alignment trimming software for accurate phylogenomic inference. PLoS Biol. 18, e3001007 (2020).

78. J. P. Rose, T. J. Kleist, S. D. Löfstrand, B. T. Drew, J. Schönenberger, K. J. Sytsma, Phylogeny, historical biogeography, and diversification of angiosperm order Ericales suggest ancient Neotropical and East Asian connections. Mol. Phylogenet. Evol. 122, 59–79 (2018).

79. M. A. Koch, B. Haubold, T. Mitchell-Olds, Comparative evolutionary analysis of chalcone synthase and alcohol dehydrogenase loci in Arabidopsis, Arabis, and related genera (Brassicaceae). Mol. Biol. Evol. 17, 1483–1498 (2000).

80. A. Franzke, D. German, I. A. Al-Shehbaz, K. Mummenhoff, *Arabidopsis*family ties: molecular phylogeny and age estimates in Brassicaceae. Taxon 58, 425–437 (2009).

81. M. A. Beilstein, N. S. Nagalingum, M. D. Clements, S. R. Manchester, S. Mathews, Dated molecular phylogenies indicate a Miocene origin for Arabidopsis thaliana. Proc. Natl. Acad. Sci. U. S. A. 107, 18724–18728 (2010).

82. T. L. P. Couvreur, A. Franzke, I. A. Al-Shehbaz, F. T. Bakker, M. A. Koch, K. Mummenhoff, Molecular phylogenetics, temporal diversification, and principles of evolution in the mustard family (Brassicaceae). Mol. Biol. Evol. 27, 55–71 (2010).

83. S. Kagale, S. J. Robinson, J. Nixon, R. Xiao, T. Huebert, J. Condie, D. Kessler, W. E. Clarke, P. P. Edger, M. G. Links, A. G. Sharpe, I. A. P. Parkin, Polyploid evolution of the Brassicaceae during the Cenozoic era. Plant Cell 26, 2777–2791 (2014).

84. N. Hohmann, E. M. Wolf, M. A. Lysak, M. A. Koch, A time-calibrated road map of Brassicaceae species radiation and evolutionary history. Plant Cell 27, 2770–2784 (2015).

85. C.-H. Huang, R. Sun, Y. Hu, L. Zeng, N. Zhang, L. Cai, Q. Zhang, M. A. Koch, I. Al-Shehbaz, P. P. Edger, J. C. Pires, D.-Y. Tan, Y. Zhong, H. Ma, Resolution of Brassicaceae phylogeny using nuclear genes uncovers nested radiations and supports convergent morphological evolution. Mol. Biol. Evol. 33, 394–412 (2016).

86. S. Mohammadin, K. Peterse, S. J. van de Kerke, L. W. Chatrou, A. A. Dönmez, K. Mummenhoff, J. C. Pires, P. P. Edger, I. A. Al-Shehbaz, M. E. Schranz, Anatolian origins and diversification of Aethionema, the sister lineage of the core Brassicaceae. Am. J. Bot. 104, 1042–1054 (2017).

87. X. Guo, J. Liu, G. Hao, L. Zhang, K. Mao, X. Wang, D. Zhang, T. Ma, Q. Hu, I. Al-Shehbaz, M. A. Koch, Plastome phylogeny and early diversification of Brassicaceae. BMC Genomics 18, 176 (2017).

88. T. Mandáková, Z. Li, M. S. Barker, M. A. Lysak, Diverse genome organization following 13 independent mesopolyploid events in Brassicaceae contrasts with convergent patterns of gene retention. Plant J. 91, 3–21 (2017).

89. X.-C. Huang, D. A. German, M. A. Koch, Temporal patterns of diversification in Brassicaceae demonstrate decoupling of rate shifts and mesopolyploidization events. Ann. Bot. 125, 29–47 (2020).

90. N. Walden, D. A. German, E. M. Wolf, M. Kiefer, P. Rigault, X.-C. Huang, C. Kiefer, R. Schmickl, A. Franzke, B. Neuffer, K. Mummenhoff, M. A. Koch, Nested whole-genome duplications coincide with diversification and high morphological disparity in Brassicaceae. Nat. Commun. 11, 3795 (2020).

91. K. P. Hendriks, C. Kiefer, I. A. Al-Shehbaz, C. D. Bailey, A. Hooft van Huysduynen, L. A. Nikolov, L. Nauheimer, A. R. Zuntini, D. A. German, A. Franzke, M. A. Koch, M. A. Lysak, Ó. Toro-Núñez, B. Özüdoğru, V. R. Invernón, N. Walden, O. Maurin, N. M. Hay, P. Shushkov, T. Mandáková, M. E. Schranz, M. Thulin, M. D. Windham, I. Rešetnik, S. Španiel, E. Ly, J. C. Pires, A. Harkess, B. Neuffer, R. Vogt, C. Bräuchler, H. Rainer, S. B. Janssens, M. Schmull, A. Forrest, A. Guggisberg, S. Zmarzty, B. J. Lepschi, N. Scarlett, F. W. Stauffer, I. Schönberger, P. Heenan, W. J. Baker, F. Forest, K. Mummenhoff, F. Lens, Global Brassicaceae phylogeny based on filtering of 1,000-gene dataset. Curr. Biol. 33, 4052–4068.e6 (2023).

92. Y. Zhao, R. Zhang, K.-W. Jiang, J. Qi, Y. Hu, J. Guo, R. Zhu, T. Zhang, A. N. Egan, T.-S. Yi, C.-H. Huang, H. Ma, Nuclear phylotranscriptomics and phylogenomics support numerous polyploidization events and hypotheses for the evolution of rhizobial nitrogen-fixing symbiosis in Fabaceae. Mol. Plant 14, 748–773 (2021).

93. E. J. M. Koenen, D. I. Ojeda, F. T. Bakker, J. J. Wieringa, C. Kidner, O. J. Hardy, R. T. Pennington, P. S. Herendeen, A. Bruneau, C. E. Hughes, The origin of the legumes is a complex paleopolyploid phylogenomic tangle closely associated with the Cretaceous-Paleogene (K-Pg) mass extinction event. Syst. Biol. 70, 508–526 (2021).

94. Y.-H. Li, G. Zhou, J. Ma, W. Jiang, L.-G. Jin, Z. Zhang, Y. Guo, J. Zhang, Y. Sui, L. Zheng, S.-S. Zhang, Q. Zuo, X.-H. Shi, Y.-F. Li, W.-K. Zhang, Y. Hu, G. Kong, H.-L. Hong, B. Tan, J. Song, Z.-X. Liu, Y. Wang, H. Ruan, C. K. L. Yeung, J. Liu, H. Wang, L.-J. Zhang, R.-X. Guan, K.-J. Wang, W.-B. Li, S.-Y. Chen, R.-Z. Chang, Z. Jiang, S. A. Jackson, R. Li, L.-J. Qiu, De novo assembly of soybean wild relatives for pan-genome analysis of diversity and agronomic traits. Nat. Biotechnol. 32, 1045–1052 (2014).

95. M. Y. Kim, S. Lee, K. Van, T.-H. Kim, S.-C. Jeong, I.-Y. Choi, D.-S. Kim, Y.-S. Lee, D. Park, J. Ma, W.-Y. Kim, B.-C. Kim, S. Park, K.-A. Lee, D. H. Kim, K. H. Kim, J. H. Shin, Y. E. Jang, K. D. Kim, W. X. Liu, T. Chaisan, Y. J. Kang, Y.-H. Lee, K.-H. Kim, J.-K. Moon, J. Schmutz, S. A. Jackson, J. Bhak, S.-H. Lee, Whole-genome sequencing and intensive analysis of the undomesticated soybean (Glycine soja Sieb. and Zucc.) genome. Proc. Natl. Acad. Sci. U. S. A. 107, 22032–22037 (2010).

96. Y. Zhuang, X. Wang, X. Li, J. Hu, L. Fan, J. B. Landis, S. B. Cannon, J. Grimwood, J. Schmutz, S. A. Jackson, J. J. Doyle, X. S. Zhang, D. Zhang, J. Ma, Phylogenomics of the genus Glycine sheds light on polyploid evolution and life-strategy transition. Nat. Plants 8, 233–244 (2022).

97. J. C. Villarreal A, B. J. Crandall-Stotler, M. L. Hart, D. G. Long, L. L. Forrest, Divergence times and the evolution of morphological complexity in an early land plant lineage (Marchantiopsida) with a slow molecular rate. New Phytol. 209, 1734–1746 (2016).

98. C.-H. Huang, C. Zhang, M. Liu, Y. Hu, T. Gao, J. Qi, H. Ma, Multiple polyploidization events across Asteraceae with two nested events in the early history revealed by nuclear phylogenomics. Mol. Biol. Evol. 33, 2820–2835 (2016).

100. J. R. Mandel, R. B. Dikow, C. M. Siniscalchi, R. Thapa, L. E. Watson, V. A. Funk, A fully resolved backbone phylogeny reveals numerous dispersals and explosive diversifications throughout the history of Asteraceae. Proc. Natl. Acad. Sci. U. S. A. 116, 14083–14088 (2019).

101. C. Zhang, C.-H. Huang, M. Liu, Y. Hu, J. L. Panero, F. Luebert, T. Gao, H. Ma, Phylotranscriptomic insights into Asteraceae diversity, polyploidy, and morphological innovation. J. Integr. Plant Biol. 63, 1273–1293 (2021).

102. J. L. Panero, B. S. Crozier, Macroevolutionary dynamics in the early diversification of Asteraceae. Mol. Phylogenet. Evol. 99, 116–132 (2016).

103. R. Chu, X. Xu, Z. Lu, Y. Ma, H. Cheng, S. Zhu, F. T. Bakker, M. E. Schranz, Z. Wei, Plastome-based phylogeny and biogeography of Lactuca L. (Asteraceae) support revised lettuce gene pool categories. Front. Plant Sci. 13, 978417 (2022).

103. N. Kilian, A. Sennikov, Z.-H. Wang, B. Gemeinholzer, J.-W. Zhang, Sub-Paratethyan origin and Middle to Late Miocene principal diversification of the Lactucinae (Compositae: Cichorieae) inferred from molecular phylogenetics, divergence-dating and biogeographic analysis. Taxon 66, 675–703 (2017).

104. K. Tremetsberger, B. Gemeinholzer, H. Zetzsche, S. Blackmore, N. Kilian, S. Talavera, Divergence time estimation in Cichorieae (Asteraceae) using a fossil-calibrated relaxed molecular clock. Org. Divers. Evol. 13, 1–13 (2013).

105. J. Bechteler, G. Peñaloza-Bojacá, D. Bell, J. Gordon Burleigh, S. F. McDaniel, E. Christine Davis, E. B. Sessa, A. Bippus, D. Christine Cargill, S. Chantanoarrapint, I. Draper, L. Endara, L. L. Forrest, R. Garilleti, S. W. Graham, S. Huttunen, J. J. Lazo, F. Lara, J. Larraín, L. R. Lewis, D. G. Long, D. Quandt, K. Renzaglia, A. Schäfer-Verwimp, G. E. Lee, A. M. Sierra, M. von Konrat, C. E. Zartman, M. R. Pereira, B. Goffinet, J. C. Villarreal A, Comprehensive phylogenomic time tree of bryophytes reveals deep relationships and uncovers gene incongruences in the last 500 million years of diversification. Am. J. Bot. 110, e16249 (2023).

106. B. J. Harris, J. W. Clark, D. Schrempf, G. J. Szöllősi, P. C. J. Donoghue, A. M. Hetherington, T. A. Williams, Divergent evolutionary trajectories of bryophytes and tracheophytes from a complex common ancestor of land plants. *Nat*. Ecol. Evol. 6, 1634–1643 (2022).

107. A. M. Kozlov, D. Darriba, T. Flouri, B. Morel, A. Stamatakis, RAxML-NG: a fast, scalable and user-friendly tool for maximum likelihood phylogenetic inference. Bioinformatics 35, 4453–4455 (2019).

108. D. M. Emms, S. Kelly, OrthoFinder: phylogenetic orthology inference for comparative genomics. Genome Biol. 20, 238 (2019).

109. One Thousand Plant Transcriptomes Initiative, One thousand plant transcriptomes and the phylogenomics of green plants. Nat. Commun.

110. B. Q. Minh, H. A. Schmidt, O. Chernomor, D. Schrempf, M. D. Woodhams, A. von Haeseler, R. Lanfear, IQ-TREE 2: New models and efficient methods for phylogenetic inference in the genomic era. Mol. Biol. Evol. 37, 1530–1534 (2020).

111. J. Kim, M. J. Sanderson, Penalized likelihood phylogenetic inference: bridging the parsimony-likelihood gap. Syst. Biol. 57, 665–674 (2008).

112. M. J. Sanderson, Estimating absolute rates of molecular evolution and divergence times: a penalized likelihood approach. Mol. Biol. Evol. 19, 101–109 (2002).

113. E. Paradis, Molecular dating of phylogenies by likelihood methods: a comparison of models and a new information criterion. Mol. Phylogenet. Evol. 67, 436–444 (2013).

114. S. Kumar, M. Suleski, J. M. Craig, A. E. Kasprowicz, M. Sanderford, M. Li, G. Stecher, S. B. Hedges, TimeTree 5: An Expanded Resource for Species Divergence Times. Mol. Biol. Evol. 39 (2022).

115. J. D. Buenrostro, B. Wu, U. M. Litzenburger, D. Ruff, M. L. Gonzales, M. P. Snyder, H. Y. Chang, W. J. Greenleaf, Single-cell chromatin accessibility reveals principles of regulatory variation. Nature 523, 486–490 (2015).

116. A. M. Bolger, M. Lohse, B. Usadel, Trimmomatic: a flexible trimmer for Illumina sequence data. Bioinformatics 30, 2114–2120 (2014).

117. P. S. Hosmani, M. Flores-Gonzalez, H. van de Geest, F. Maumus, L. V. Bakker, E. Schijlen, J. van Haarst, J. Cordewener, G. Sanchez-Perez, S. Peters, Z. Fei, J. J. Giovannoni, L. A. Mueller, S. Saha, An improved de novo assembly and annotation of the tomato reference genome using single-molecule sequencing, Hi-C proximity ligation and optical maps, Genomics (2019). https://www.biorxiv.org/content/10.1101/767764v1.abstract.

118. H. Li, Aligning sequence reads, clone sequences and assembly contigs with BWA-MEM, *arXiv [q-bio.GN]* (2013). http://arxiv.org/abs/1303.3997.

119. H. Zhao, Z. Sun, J. Wang, H. Huang, J.-P. Kocher, L. Wang, CrossMap: a versatile tool for coordinate conversion between genome assemblies. Bioinformatics 30, 1006–1007 (2014).

120. X. Tu, M. K. Mejía-Guerra, J. A. Valdes Franco, D. Tzeng, P.-Y. Chu, W. Shen, Y. Wei, X. Dai, P. Li, E. S. Buckler, S. Zhong, Reconstructing the maize leaf regulatory network using ChIP-seq data of 104 transcription factors. Nat. Commun. 11, 5089 (2020).

121. M. Galli, A. Khakhar, Z. Lu, Z. Chen, S. Sen, T. Joshi, J. L. Nemhauser, R. J. Schmitz, A. Gallavotti, The DNA binding landscape of the maize AUXIN RESPONSE FACTOR family. Nat. Commun. 9, 4526 (2018).

122. M. Galli, Z. Chen, T. Ghandour, A. Chaudhry, J. Gregory, F. Feng, M. Li, N. Schleif, X. Zhang, Y. Dong, G. Song, J. W. Walley, G. Chuck, C. Whipple, H. F. Kaeppler, S.-S. C. Huang, A. Gallavotti, Transcription factor binding divergence drives transcriptional and phenotypic variation in maize. Nat. Plants 11, 1205–1219 (2025).

123. W. J. Kent, A. S. Zweig, G. Barber, A. S. Hinrichs, D. Karolchik, BigWig and BigBed: enabling browsing of large distributed datasets. Bioinformatics 26, 2204–2207 (2010).

124. G. Ambrosini, R. Groux, P. Bucher, PWMScan: a fast tool for scanning entire genomes with a position-specific weight matrix. Bioinformatics 34, 2483–2484 (2018).

125. Z. Lu, B. T. Hofmeister, C. Vollmers, R. M. DuBois, R. J. Schmitz, Combining ATAC-seq with nuclei sorting for discovery of cis-regulatory regions in plant genomes. Nucleic Acids Res. 45, e41 (2017).

126. C. M. Alexandre, J. R. Urton, K. Jean-Baptiste, J. Huddleston, M. W. Dorrity, J. T. Cuperus, A. M. Sullivan, F. Bemm, D. Jolic, A. A. Arsovski, A. Thompson, J. L. Nemhauser, S. Fields, D. Weigel, K. L. Bubb, C. Queitsch, Complex relationships between chromatin accessibility, sequence divergence, and gene expression in Arabidopsis thaliana. Mol. Biol. Evol. 35, 837–854 (2018).

127. Y. Liu, T. Tian, K. Zhang, Q. You, H. Yan, N. Zhao, X. Yi, W. Xu, Z. Su, PCSD: a plant chromatin state database. Nucleic Acids Res. 46, D1157–D1167 (2018).

128. Y. Zhang, T. Liu, C. A. Meyer, J. Eeckhoute, D. S. Johnson, B. E. Bernstein, C. Nusbaum, R. M. Myers, M. Brown, W. Li, X. S. Liu, Model-based analysis of ChIP-Seq (MACS). Genome Biol. 9, R137 (2008).

129. F. Hammal, P. de Langen, A. Bergon, F. Lopez, B. Ballester, ReMap 2022: a database of Human, Mouse, Drosophila and Arabidopsis regulatory regions from an integrative analysis of DNA-binding sequencing experiments. Nucleic Acids Res. 50, D316–D325 (2022).

130. H. Stroud, M. V. C. Greenberg, S. Feng, Y. V. Bernatavichute, S. E. Jacobsen, Comprehensive analysis of silencing mutants reveals complex regulation of the Arabidopsis methylome. Cell 152, 352–364 (2013).

131. J. An, R. Brik Chaouche, L. I. Pereyra-Bistraín, H. Zalzalé, Q. Wang, Y. Huang, X. He, C. Dias Lopes, J. Antunez-Sanchez, C. Bergounioux, C. Boulogne, C. Dupas, C. Gillet, J. M. Pérez-Pérez, O. Mathieu, N. Bouché, S. Fragkostefanakis, Y. Zhang, S. Zheng, M. Crespi, M. M. Mahfouz, F. Ariel, J. Gutierrez-Marcos, C. Raynaud, D. Latrasse, M. Benhamed, An atlas of the tomato epigenome reveals that KRYPTONITE shapes TAD-like boundaries through the control of H3K9ac distribution. Proc. Natl. Acad. Sci. U. S. A. 121, e2400737121 (2024).

132. H. Patel, J. Espinosa-Carrasco, C. Wang, P. Ewels, N.-C. Bot, T. C. Silva, A. Peltzer, B. Langer, S. Guinchard, M. U. Garcia, D. Behrens, M. Hörtenhuber, A. Talbot, K. Rokicki, R. Syme, Rotholandus, S. R. Pérez, S. Haglund, S. Möller, W. W. “winni” Kretzschmar, K. Menden, Nf-Core/chipseq: Nf-Core/chipseq v2.1.0 - Platinum Willow Sparrow (Zenodo, 2024; 10.5281/ZENODO.3240506).

133. F. Krueger, F. James, P. Ewels, E. Afyounian, M. Weinstein, B. Schuster-Boeckler, G. Hulselmans, sclamons, FelixKrueger/TrimGalore: v0.6.10 - Add Default Decompression Path (Zenodo, 2023; 10.5281/zenodo.7598955).

134. *Picard: A Set of Command Line Tools (in Java) for Manipulating High-Throughput Sequencing (HTS) Data and Formats such as SAM/BAM/CRAM and VCF* (Github; https://github.com/broadinstitute/picard).

135. H. Li, B. Handsaker, A. Wysoker, T. Fennell, J. Ruan, N. Homer, G. Marth, G. Abecasis, R. Durbin, 1000 Genome Project Data Processing Subgroup, The Sequence Alignment/Map format and SAMtools. Bioinformatics 25, 2078–2079 (2009).

136. D. W. Barnett, E. K. Garrison, A. R. Quinlan, M. P. Strömberg, G. T. Marth, BamTools: a C++ API and toolkit for analyzing and managing BAM files. Bioinformatics 27, 1691–1692 (2011).

137. A. R. Quinlan, I. M. Hall, BEDTools: a flexible suite of utilities for comparing genomic features. Bioinformatics 26, 841–842 (2010).

138. G. Yu, L.-G. Wang, Y. Han, Q.-Y. He, clusterProfiler: an R package for comparing biological themes among gene clusters. OMICS 16, 284–287 (2012).

139. AnnotationHub, *Bioconductor*. https://bioconductor.org/packages/AnnotationHub.

140. J. Van Eck, P. Keen, M. Tjahjadi, Agrobacterium tumefaciens-mediated transformation of tomato. Methods Mol. Biol. 1864, 225–234 (2019).

141. D. Rodríguez-Leal, Z. H. Lemmon, J. Man, M. E. Bartlett, Z. B. Lippman, Engineering quantitative trait variation for crop improvement by genome editing. Cell 171, 470–480.e8 (2017).

142. S. Soyk, Z. H. Lemmon, M. Oved, J. Fisher, K. L. Liberatore, S. J. Park, A. Goren, K. Jiang, A. Ramos, E. van der Knaap, J. Van Eck, D. Zamir, Y. Eshed, Z. B. Lippman, Bypassing negative epistasis on yield in tomato imposed by a domestication gene. Cell 169, 1142–1155.e12 (2017).

143. S. Werner, C. Engler, E. Weber, R. Gruetzner, S. Marillonnet, Fast track assembly of multigene constructs using Golden Gate cloning and the MoClo system. Bioeng. Bugs 3, 38–43 (2012).

144. D. T. Hoang, O. Chernomor, A. von Haeseler, B. Q. Minh, L. S. Vinh, UFBoot2: Improving the ultrafast bootstrap approximation. Mol. Biol. Evol. 35, 518–522 (2018).

145. G. Yu, D. K. Smith, H. Zhu, Y. Guan, T. T.-Y. Lam, Ggtree: An r package for visualization and annotation of phylogenetic trees with their covariates and other associated data. Methods Ecol. Evol. 8, 28–36 (2017).

146. L.-G. Wang, T. T.-Y. Lam, S. Xu, Z. Dai, L. Zhou, T. Feng, P. Guo, C. W. Dunn, B. R. Jones, T. Bradley, H. Zhu, Y. Guan, Y. Jiang, G. Yu, Treeio: An R package for phylogenetic tree input and output with richly annotated and associated data. Mol. Biol. Evol. 37, 599–603 (2020).

147. T. Hackl, M. Ankenbrand, B. van Adrichem, D. Wilkins, K. Haslinger, gggenomes: effective and versatile visualizations for comparative genomics, arXiv [q-bio.GN] (2024). http://arxiv.org/abs/2411.13556.

148. P. Danecek, J. K. Bonfield, J. Liddle, J. Marshall, V. Ohan, M. O. Pollard, A. Whitwham, T. Keane, S. A. McCarthy, R. M. Davies, H. Li, Twelve years of SAMtools and BCFtools. Gigascience 10 (2021).

149. A. M. Waterhouse, J. B. Procter, D. M. A. Martin, M. Clamp, G. J. Barton, Jalview Version 2--a multiple sequence alignment editor and analysis workbench. Bioinformatics 25, 1189–1191 (2009).

150. Z. B. Lippman, O. Cohen, J. P. Alvarez, M. Abu-Abied, I. Pekker, I. Paran, Y. Eshed, D. Zamir, The making of a compound inflorescence in tomato and related nightshades. PLoS Biol. 6, e288 (2008).

151. N. Servant, N. Varoquaux, B. R. Lajoie, E. Viara, C.-J. Chen, J.-P. Vert, E. Heard, J. Dekker, E. Barillot, HiC-Pro: an optimized and flexible pipeline for Hi-C data processing. Genome Biol. 16, 259 (2015).

152. M. Alonge, L. Lebeigle, M. Kirsche, K. Jenike, S. Ou, S. Aganezov, X. Wang, Z. B. Lippman, M. C. Schatz, S. Soyk, Automated assembly scaffolding using RagTag elevates a new tomato system for high-throughput genome editing. Genome Biol. 23, 258 (2022).

153. T. Yang, F. Zhang, G. G. Yardımcı, F. Song, R. C. Hardison, W. S. Noble, F. Yue, Q. Li, HiCRep: assessing the reproducibility of Hi-C data using a stratum-adjusted correlation coefficient. Genome Res. 27, 1939–1949 (2017).

154. N. Abdennur, L. A. Mirny, Cooler: scalable storage for Hi-C data and other genomically labeled arrays. Bioinformatics 36, 311–316 (2020).

155. A. Kaul, S. Bhattacharyya, F. Ay, Identifying statistically significant chromatin contacts from Hi-C data with FitHiC2. Nat. Protoc. 15, 991–1012 (2020).

156. H. Li, Minimap2: pairwise alignment for nucleotide sequences. Bioinformatics 34, 3094–3100 (2018).

157. K. A. Riemondy, R. M. Sheridan, A. Gillen, Y. Yu, C. G. Bennett, J.R. Hesselberth, valr: Reproducible genome interval analysis in R. F1000Res. 6, 1025 (2017).

158. X. Qiao, Q. Li, H. Yin, K. Qi, L. Li, R. Wang, S. Zhang, A. H. Paterson, Gene duplication and evolution in recurring polyploidization-diploidization cycles in plants. Genome Biol. 20, 38 (2019).

159. N. Uchida, B. Townsley, K.-H. Chung, N. Sinha, Regulation of SHOOT MERISTEMLESS genes via an upstream-conserved noncoding sequence coordinates leaf development. Proc. Natl. Acad. Sci. U. S. A. 104, 15953–15958 (2007).

